# Pure Nano Genesis: Pioneering Universal Aqueous Nanostrategies from Pure Molecules to Revolutionise Diverse Applications

**DOI:** 10.1101/2024.03.24.586367

**Authors:** Wenwen Zhao, Qiu Li, Peng He, Changqing Li, Muna Aryal, Mario L. Fabiilli, Haijun Xiao

## Abstract

In this paper, we propose a novel universal approach for the construction of stable and aqueous nanoparticles, Pure Nano systems, comprising solely small conjugated molecules without any excipients. Our experiments confirm that the generation of surface charges plays an essential role during the spontaneous aggregation of conjugated molecules in the process of Pure Nano system fabrication, as it governs growth and confers physiochemical stability to particles at the nano scale. This approach overcomes solubility challenges in highly hydrophobic conjugated molecules by obviating excipients and enabling up to 100% drug loading capacity. Confirmation of this capability stems from the successful preparation of approximately 100 Pure Nano systems, incorporating different combinations of 27 conjugated molecules distinguished by their diverse dissociation types and degrees. The proposed preparation method is robust, simple, fast, and reliable, making it well-suited for large-scale manufacturing due to its array of unique features. This strategy affords a singular, molecular-focused approach, showcasing the intrinsic bioactivity of its constituent molecules while enabling aqueous dispersion for diverse applications. And *in vivo* experiments confirm the exceptional efficacy of various Pure Nano systems in reinstating dextran sodium sulfate induced acute ulcerative colitis to a healthy state, restoring myocardial ischemia/reperfusion injury to normal levels, and effectively treating cancer in mice with significantly improved median survival rates. This innovative nano drug delivery system represents a groundbreaking advancement with the potential to revolutionise translational nanomedicine. Positioned at the forefront of therapeutic possibilities, it is anticipated to substantially improve the efficacy and safety of nano therapies. This advancement heralds a new era, offering a highly targeted and efficient approach in the treatment of diverse medical conditions.

## 1 Introduction

Nanoparticles have garnered significant attention as drug delivery systems due to their distinct features, including enhanced drug bioavailability and reduced adverse effects, thereby facilitating the creation of more effective and safe therapeutics.^1^ However, despite decades of advancements, some challenges continue to persist in this field.

Conventional approaches to nanoparticle generation are heavily reliant on excipients, with polymers being a prominent category employed in nanoparticle preparation.^2,3^ The formation strategy revolves around exploiting the amphiphilic traits of the excipients, empowering their self-assembly into nano structures.^4,5^ The excipients are originally incorporated into the drug delivery systems to address the limited water solubility of highly hydrophobic drug molecules^3^, but in turn, can create additional complications.

Poor stability poses a significant obstacle to the clinical translation of numerous nanoparticle systems. A fundamental reason for this impediment lies in the inherently weak intermolecular interactions, such as Van der Waals forces (London dispersion force and dipole-dipole forces), hydrogen bonding and hydrophobic interactions (Tendency of hydrophobic moieties to aggregate in aqueous solution), that exist among the assembly chains constituting the nanoparticle skeleton.^6–8^ Accordingly, nanoparticles formulated *via* conventional amphiphilic components are particularly vulnerable to instability, thereby engendering phenomena such as coalescence or flocculation, prior to their successful arrival at targeted sites.^8,9^ Furthermore, the complex *in vivo* milieu further exacerbates the stability concerns that excipient-based nanoparticles encounter, primarily *via* interactions with biological constituents, as well as dilution in the blood.^10^ Successful drug delivery entails improved stability of nanoparticles, and hence demands feasible strategies to optimise this feature. In pursuit of this goal, diverse approaches, including cross-linking, hybridisation and covalent modification, have been probed to bolster the structural stability of nano-formulations.^11–13^

Another pivotal impediment to the efficacy of excipient-based nanoparticles is their inadequate drug loading capacity, which undermines therapeutic effectiveness and consequently compromises clinical outcomes.^14–16^ Additionally, suboptimal drug loading can engender the need for increased dosing frequencies or concentrative surpluses, culminating in patient inconvenience, escalated healthcare expenditures and compromised patient compliance. In the industrial context, suboptimal drug payload may result in heightened production costs due to the necessity for greater quantities of excipients. And the production of auxiliary materials involves additional investments of labour, resources and time. These may consequently constrain the commercial feasibility of drug delivery systems. Furthermore, the scarcity of products endowed with high loading capacities for drugs further compounds this issue.

In the realm of nanotechnology for *in vivo* use, only a restricted selection of polymers, such as poly(lactic-co-glycolic acid) (PLGA) and polyethylene glycol (PEG), have obtained approval for applications so far.^17,18^ PEG, lauded for its favourable safety profile and good water solubility, has garnered significant employment in a multitude of products and investigations. Nonetheless, it poses a notable risk of provoking severe allergic responses in some patients.^19–21^ Other materials that have not been validated over time for use *in vivo* will pose additional safety concerns, especially if they accumulate in organs over time.

As a result, alternative strategies are essential to overcoming the multifaceted issues associated with the conventional technique for nanoparticle construction to facilitate the development of superior drug delivery systems. Novel approaches that offer enhanced control over particle size, increased drug loading capacity and biocompatibility, could improve current limitations and extend the scope of nanoparticle drug delivery.

Here, we report a novel universal approach for the construction of stable and aqueous nanoparticles, Pure Nano systems, comprising solely small conjugated molecules inherently endowed with bioactive attributes, absent of supplementary excipients or adjunct compounds. The spontaneous aggregation of conjugated molecules during Pure Nano system fabrication hinges upon the pivotal role of surface charge generation. This process distinctly governs particle growth and bestows crucial physiochemical stability at the nano scale. This approach emphasises a molecular-centric strategy, highlighting the intrinsic bioactivity of its constituent molecules and enabling versatile aqueous dispersion for diverse applications. Notably, *in vivo* experiments corroborate the exceptional efficacy of various Pure Nano systems. These systems exhibit promising results in ameliorating Dextran Sodium Sulfate (DSS)-induced acute ulcerative colitis, mitigating myocardial ischemia/reperfusion (I/R) injury, and demonstrating efficacy in mouse models of cancer. Additionally, these systems demonstrate potential applications in NIR fluorescence imaging, X-ray-based imaging, and various forms of photo-based therapy such as photothermal and photodynamic therapies. This discovery has been shown to be a universal approach with the potential to revolutionise nano drug delivery systems by enabling the development of stable and aqueous nanoparticles without the use of excipients, which offers the capability to address the limitations of conventional drug delivery strategies and facilitate the development of safer and more effective nano therapies.

## 2 Materials and Methods

The Pure Nano systems were produced utilising a quick and straightforward self aggregation and dispersion method with some modifications^22^. In brief, a certain amount of small molecules (listed in Table 1) with specific ratios were dissolved in organic solvents to produce a combined solution, which was then precipitated in an aqueous solution to generate a nanoparticle dispersion. The detailed specifications for each formulation was displayed in Table 2 and 3, encompassing the combinations of molecules (molecular grouping), mass quantities, molecular ratios, the designated organic solvent as the oil phase, *pH* levels of the aqueous phase, and the respective volumes of the aqueous phase. The nanoparticle powder was obtained through the meticulous process of solvent removal using lyophilisation, a technique involving freeze-drying to eliminate the solvent content. This method ensured the preservation of the nanoparticle’s structural integrity and stability for subsequent applications. The resulting product was stored under light-protected conditions at 4°C to safeguard against degradation until further utilisation.

**Table 1:** Molecular Identification Numbers and Physicochemical Properties.

| Molecule No. | Name | Category<sup>1</sup> | CAS No. | Mass (g/mol) | LogP<sup>2</sup> | [S]<sup>3</sup> | pI<sup>4</sup> | pK<sub>a</sub> |
| --- | --- | --- | --- | --- | --- | --- | --- | --- |
| 1 | Psoralen | NI | 66-97-7 | 186 | 2.13 | 0.24 | — | — |
| 2 | Pyrene | NI | 129-00-0 | 202 | 5.19 | 0.00 | — | — |
| 3 | Tryptanthrin | NI | 13220-57-0 | 248 | 2.31 | 0.35 | — | — |
| 4 | Tanshinone I | NI | 568-73-0 | 276 | 4.10 | 0.01 | — | — |
| 5 | Pigment violet 23 | NI | 215247-95-3 | 589 | 7.53 | 0.01 | — | — |
| 6 | Aristolochic acid | A | 313-67-7 | 341 | 2.69 | 0.03 | — | 3.15 |
| 7 | Rhein | A | 478-43-3 | 284 | 2.18 | 0.22 | — | 3.40 |
| 8 | Ellagic acid | A | 476-66-4 | 302 | 1.59 | 0.81 | — | 5.54 |
| 9 | Hypericin | A | 548-04-9 | 504 | 3.92 | 0.01 | 2.62 | 6.65 |
| 10 | Naphthofluorescein | A | 61419-02-1 | 432 | 4.97 | 0.00 | — | 9.45 |
| 11 | Tanshinon IIB | A | 17397-93-2 | 310 | 3.31 | 0.03 | — | 14.98 |
| 12 | Camptothecin | B | 7689-03-4 | 348 | 1.91 | 0.52 | 7.40 | 3.07 |
| 13 | Nile Red | B | 7385-67-3 | 318 | 4.39 | 0.03 | — | 3.90 |
| 14 | Ellipticine | B | 519-23-3 | 246 | 4.34 | 0.00 | 10.97 | 5.12 |
| 15 | Amonafide | B | 69408-81-7 | 283 | 0.90 | 0.76 | — | 8.52 |
| 16 | Anonaine | B | 1862-41-5 | 265 | 2.22 | 0.03 | — | 9.20 |
| 17 | Mitoxantrone dihydrochloride | CA | 70476-82-3 | 517 | 0.91 | 89.00 | 9.03 | 8.86 |
| 18 | Topotecan hydrochloride | CA | 119413-54-6 | 458 | 1.84 | 92.00 | 8.94 | 9.89 |
| 19 | Sodium Tanshinone IIA Sulfonate | CB | 69659-80-9 | 398 | 3.34 | 10.00 | — | -2.41 |
| 20 | Acid Green 25 | CB | 4403-90-1 | 623 | 4.09 | 10.00 | — | -1.93 |
| 21 | IR-820 | CB | 172616-80-7 | 849 | 4.34 | 0.00 | — | — |
| 22 | Rose bengal sodium | CB | 632-69-9 | 1018 | 5.84 | 25.00 | — | 2.38 |
| 23 | Eosin Y disodium salt | CB | 17372-87-1 | 692 | 6.12 | 40.00 | — | 3.37 |
| 24 | Methylene blue | PC, B | 61-73-4 | 320 | -1.08 | 43.60 | — | 2.44 |
| 25 | Rhodamine B | PC, A | 81-88-9 | 479 | 2.37 | 50.00 | — | 3.50 |
| 26 | Berberine chloride | PC | 141433-60-5 | 372 | -0.18 | 1.95 | — | — |
| 27 | Curcumin | A | 458-37-7 | 368 | 3.62 | 0.00 | — | 7.80 |

*Notes:*
LogP, solubility and pK<sub>a</sub> values were sourced from public databases such as ChemSpide, PubChem and DrugBank, or predicted using software tools such as ALOGPS and JChem. The reported values for a particular property can vary slightly between different sources due to factors such as the method of measurement, experimental conditions and/or calculation protocols used.
<sup>1</sup> NI, Non-Ionisable molecules; A, Acids; B, Bases; CA, Conjugate Acids of a base; CB, Conjugate Bases of an acid; PC, Permanently Charged molecules.
<sup>2</sup> LogP > 0, molecules are hydrophobic; LogP < 0, molecules are hydrophilic.
<sup>3</sup> [S]: Solubility in water at room temperature and pressure (mg/mL). [S] < 1 mg/mL, molecules are very slightly soluble or practically insoluble in water.
<sup>4</sup> pI is short for isoelectric point.

**Table 2:**
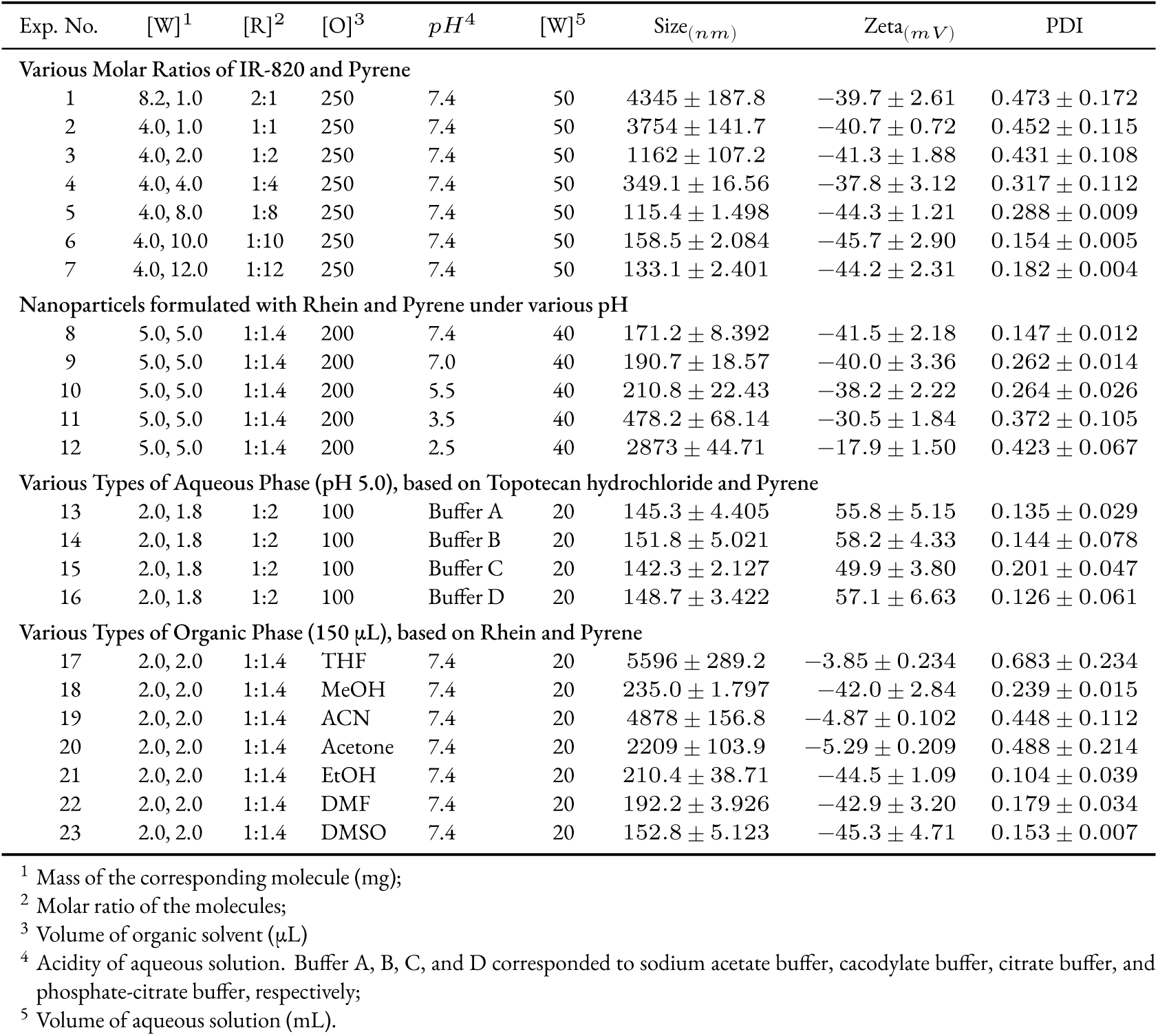
Effects of formulation parameters on nanoparticle formation.

The comprehensive details regarding the formulation and preparation of these nano systems, along with materials and methods for both *in vitro* and *in vivo* experiments, are provided in the Supplementary Information for thorough information.

## 3 Results and Discussion

### 3.1 Molecular Properties

To investigate the spontaneous dissociation and aggregation behaviour, the physicochemical characteristics of 27 diverse chemical molecules were assessed (Table 1). The molecular weights of the compounds span a wide range, from around 200 g/mol to over 1000 g/mol, correlating with varied biological activities such as enzyme inhibition and receptor binding. According to the solubility criteria of United States Pharmacopeia (USP)^23^, the molecules exhibit extremely limited solubility in water under room temperature and pressure, with values typically less than 1 mg/mL, except for those that can form salts.

#### 3.1.1 Spontaneous Dissociation of Molecules in Aqueous Phase

The molecules in Table 1 can be classified into different categories based on their dissociation types and degrees in the field of acid-base dissociation. Ionisable molecules are those that can dissociate into ions in aqueous solution and are further classified as acidic or basic based on their ability to donate or accept protons (H^+^). Acidic molecules typically contain ionisable acidic groups or both ionisable acidic and basic groups, with the latter having an isoelectric point of less than 7. Basic compounds, on the other hand, consist of either ionisable basic groups or both ionisable acidic and basic groups, provided that their isoelectric point is greater than 7.^24^ The conjugate acids of a base are the species that form when the base accepts one or more protons, while the conjugate bases of an acid result after the acid donates one or more protons.

Furthermore, molecules that are permanently charged are those that maintain their charges regardless of the *pH* of the surrounding environment. Permanently negatively charged molecules are relatively rare. In contrast, there are several types of molecules that are permanently positively charged, such as quaternary ammonium compounds (R_4_N^+^), sulfonium compounds (R_3_S^+^), phosphonium compounds (R_3_P^+^), imidazolium compounds (R_2_C_3_N_2_H_3_^+^) and pyridinium compounds (C_5_H_5_NH^+^).^25–28^ These molecules contain positively charged functional groups that are not affected by changes in *pH* because their charges are delocalised over multiple atoms or are stabilised by resonance effects.

The *pK_a_* value is a fundamental measure of acid strength that evaluates the proton affinity of an acid in the context of Bronsted acid-base theory.^29^ A lower *pK_a_* value signifies a stronger acid with a greater tendency to donate protons. On the other hand, for a base, the strength can be ascertained by the *pK_a_* value of its conjugate acid. In general, a lower *pK_a_* of the conjugate acid indicates a stronger acid and, consequently, a weaker base.

The Henderson-Hasselbalch equation describes the relationship between the *pH*, *pK_a_* and the disassociation of a weak acid *HA* or a conjugate acid of a weak base *B* in an aqueous solution.^30^ The equation for the dissociation of a weak acid is:

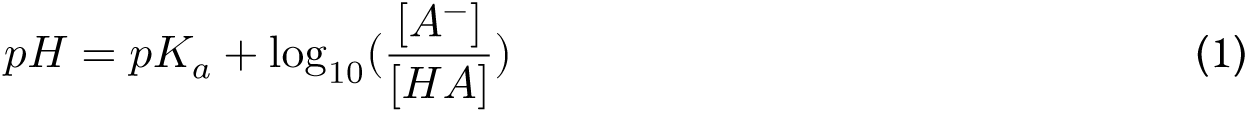

and the equation for the dissociation of a conjugate acid of a weak base is:

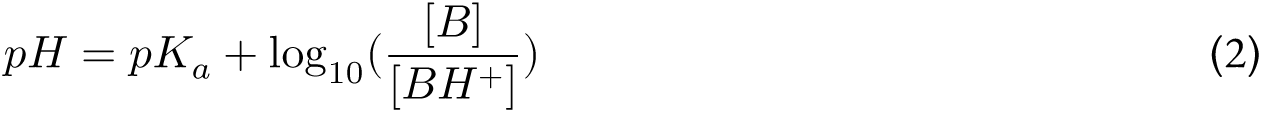

Here, *pH* represents the acidity of an aqueous solution, *K*_*a*_ is the disassociation constant of a weak acid or a conjugate acid of a weak base. *pK_a_* is the negative logarithm of *K*_*a*_ values. At equilibrium, the concentration of a weak acid is denoted as [*HA*] and the concentration of its conjugate base is denoted as [*A*^−^]. Similarly, the concentration of a weak base at equilibrium is represented by [*B*] and the concentration of its conjugate acid is denoted as [*BH*^+^].

The Henderson-Hasselbalch equation is commonly used to understand and control the *pH* of a solution, as well as to calculate the concentration of the acid or base forms of weak acids or bases. Thus, the percentage content of molecules in their ionic states can also be expressed by the following equations, respectively:

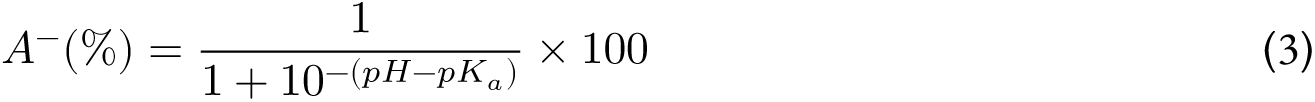

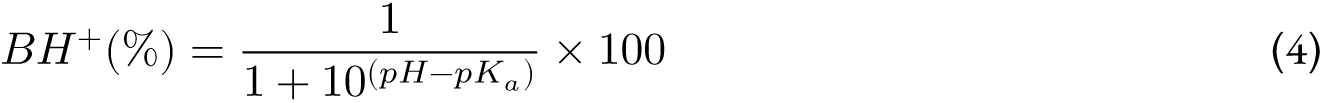

Figure 6A illustrates the correlation between the percentage of ionised molecules and the discrepancy between the *pH* of an aqueous solution and the *pK_a_* of the weak acid or base in an aqueous solution. As can be seen from the plot, the ionisation percentage of the weak acid or base is dependent on the *pH* of the solution and the *pK_a_* of the weak acid or base, while the non-ionisable molecules and permanently charged molecules remain the same regardless of changes in environmental acidity.

At the *pK_a_* value of a weak acid or base (*pH* = *pK_a_*), half of the molecules will dissociate into their ionic states, resulting in a 50% concentration of ions. In an aqueous solution with a *pH* two units higher than the *pK_a_* value of a weak acid, approximately 99% of molecules will be in an ionic state, and mainly negatively charged. Conversely, in an aqueous solution with a *pH* two units lower than the *pK_a_* value of a weak base, more than 99% of molecules will be in an ionic state, but with positive charges. The pale lavender shade represents the positively charged area under specified environmental *pH* ranges, while the lime sorbet shade represents the negatively charged area.

#### 3.1.2 Spontaneous Aggregation of Conjugated Molecules in Aqueous Phase

The chemical structures corresponding to the molecules in Table 1 are depicted in Figure 1A. These structures exhibit numerous conjugated moieties, characterized by interconnected p-orbitals with delocalized electrons^31^. This structure enables the formation of various forms of conjugation systems, such as *π* − *π* conjugation, *p* − *π* conjugation, cross conjugation and *σ* − *π* hyper conjugation.^32–35^ These conjugations are typically denoted through the alternation of single and multiple bonds, as well as the possible presence of lone pairs, radicals, or carbenium ions.^36–39^

**Figure 1:**
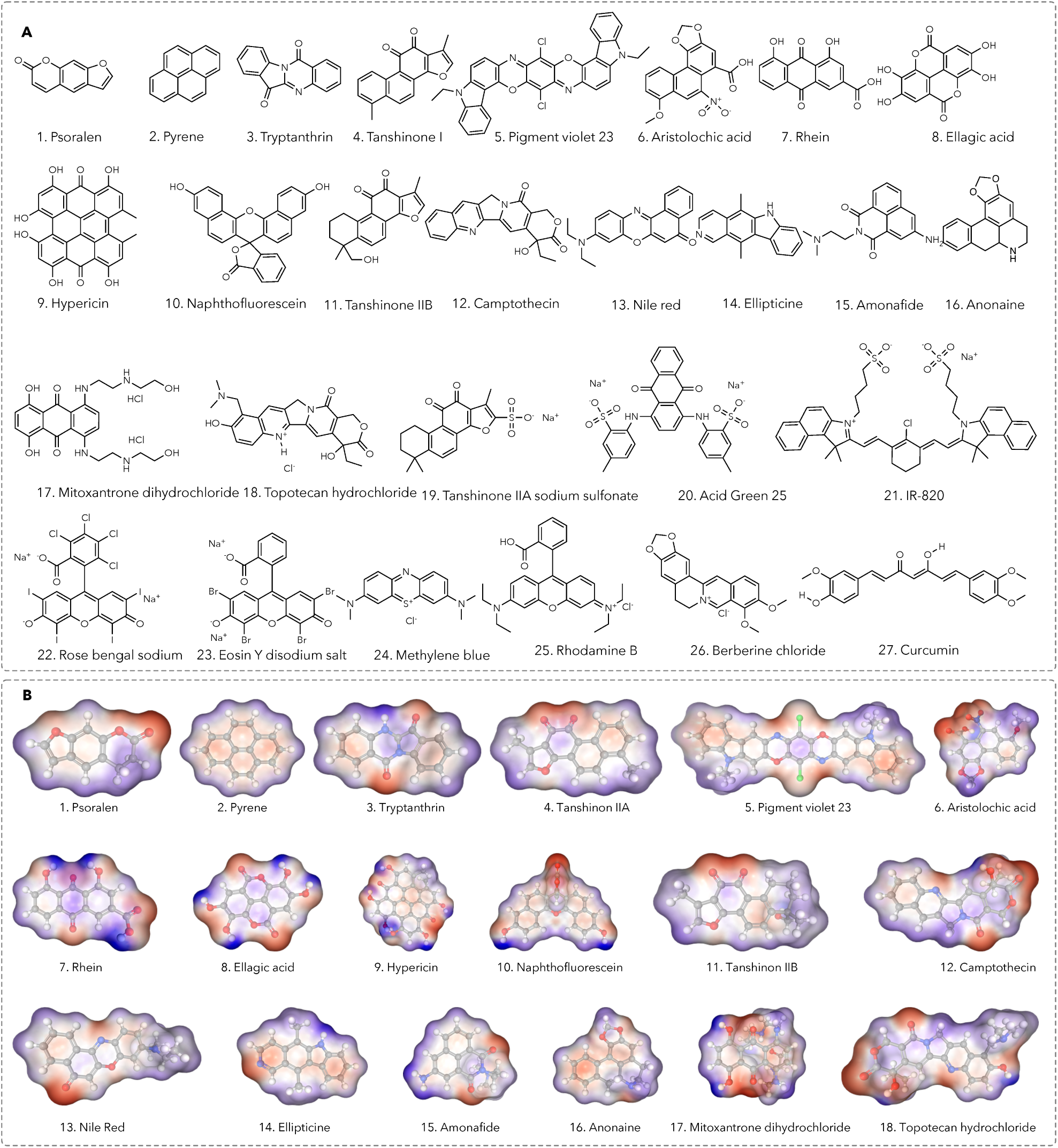
(A) The chemical structures of conjugated molecules; (B) The electrostatic potential surfaces (ESP) of molecules predicted using Graph-Convolutional Deep Neural Network **(DNN),** the colour gradient depicts the electrostatic potential distribution in molecules, which reflects the potential energy associated with the distribution of electric charge. The gradient spans the negative-to-positive range, with negative potentials indicated by shades of red and positive potentials by shades of blue.

Besides, as shown in the electrostatic potential surface (EPS, Figure lB), oxygen atoms, characterized by their high electronegativity, appear as a dark red color, signifying their strong electron-attracting nature in a chemical bond^40^. Conversely, hydrogen atoms directly bonded to oxygen exhibit a positive electrostatic potential due to their comparatively weaker electron-attracting ability. Thus, due to the conjugated system and the atoms with high electronegativities, the overall electron cloud of the molecule is unevenly distributed, forming electron-rich and electron-poor regions within molecules, which in turn leads to the difference in charges.^41^

The process of molecule aggregation is attributed to the intermolecular forces resulting from the *π* − *π* stacking interactions between the conjugated systems, as shown in 5A and B. The overlap of connected p-orbitals in adjacent conjugated systems engenders interaction between their electron clouds, producing Van der Waals forces of attraction that enable the formation of a continuous network of delocalised electrons.^42^ Besides, the difference in charges within a conjugated system can further facilitate the aggregation of conjugated systems due to the electrostatic attraction caused by the opposite charges. The negatively charged sections of a conjugated molecule would be further attracted by the positively charged regions of another.

In addition, as shown in the table 1, the molecules used in this study with logP values greater than one indicate that they are more hydrophobic in nature. This hydrophobic nature causes them to preferentially aggregate with each other in an aqueous phase, minimising their contact with water molecules.^43,43^ As a result, in uncontrolled environments, the spontaneous aggregation of molecules can result in significantly large particles with increased surface area.

### 3.2 Pure Nano Preparation and Optimisation

Pure Nano, an aqueous nanoparticle formulation comprising solely small chemical molecules was achieved by leveraging the spontaneous dissociation and aggregation behaviour of conjugated molecules.^22^ To explore significant factors influencing the nano formulation, optimisation of the Pure Nano systems involved studying various formulation parameters (molecular groupings, constituent molecule molar ratios, organic solvents, aqueous solutions) and preparative conditions (temperature, time, stirring speed, mutual addition of organic and aqueous phases).

#### 3.2.1 Formulation Optimisation

As shown in Table 2, variations in the molar ratios of components during the preparation of nanoparticles, based on IR-820 and pyrene, resulted in significant alterations in particle sizes, distributions (PDI) and zeta potentials. By modifying the ratio of IR-820 and pyrene from 2:1 to 1:12, while keeping all other formation and preparative parameters constant, a significant reduction in particle size from approximately 4000 *nm* to 100 *nm* was achieved. The enlarged conjugated area of IR-820 relative to pyrene may explain the observed effect, as the greater proportion of pyrene increases the stacked surface area. Additionally, a narrow size distribution with a PDI value of around 0.1 was observed, indicating excellent uniformity and homogeneous dispersion within the solvents. It can be inferred from these results that the optimisation of particle sizes and their distribution in this nanoparticle system can be achieved *via* appropriate adjustment of the component ratio. Moreover, the obtained nanoparticles exhibited excellent stability as evidenced by a very high zeta potential value of around −40 *mV*.

The effects of aqueous acidity on nanoparticle formation were investigated for nanoparticle preparation utilising rhein and pyrene. It was found that increasing the environmental acidity of the aqueous solution resulted in an increase in particle sizes from nano to micro sizes with a considerably wider distribution. In addition, the surface charges of the particles were also observed to decrease from around −40 *mV* to about −10 *mV* . The decrease in surface charges of the particles upon increasing the environmental acidity lends further support to the notion that the nanoparticle formation process utilising rhein and pyrene is highly sensitive to changes in aqueous acidity.

In view of the crucial role played by aqueous acidity in nanoparticle formation, the influence of buffer salts on nanoparticle formation was also explored by employing topotecan hydrochloride and pyrene. Buffer salts exhibit varied buffer ranges, yet in a suitable range with adequate buffer capacity, their impact on nanoparticle formation is negligible. Therefore, it can be inferred that the buffer salt choice in nanoparticle preparation utilising topotecan hydrochloride and pyrene may not be a crucial factor under these conditions.

The effects of organic solvents on nanoparticle formation were also explored in the context of nanoparticle preparation based on rhein and pyrene. Varying the organic solvents led to a dramatic variation in particle sizes, ranging from nano to micro sizes. Furthermore, the surface charges of the particles varied significantly, shifting from around −5 *mV* to approximately −40 *mV* . These observations are probably attributed to the different physiochemical properties of the solvents, such as their polarity, proticity, and their capacity to dissolve the molecules in the solvents.

#### 3.2.2 Preparative Parameters

Furthermore, several preparative parameters, such as temperature, stirring speed, time, and mutual addition of organic and aqueous phases during the preparation process, were also studied for their influence on nanoparticle formation. For example, varying the environmental temperature from room temperature to 0°C or changing the order of mutual addition of organic and aqueous solvents, as well as increasing the mixing time from 5 to 30 minutes, had little effects on the size and surface charges of the particles generated.

The contribution of preparative parameters to nanoparticle formation is comparatively negligible in comparison to the significant impact of formulation parameters, such as molar ratios, organic solvents, and aqueous solutions. The formulation parameters, such as molecular groupings, molar ratios of constituent molecules, and *pH* of aqueous solutions, are the key drivers of nanoparticle formation in these nanoparticle systems. This also implies that the preparation method utilised in this system is robust, simple, fast and reliable. Collectively, these findings offer crucial insights into the factors regulating nanoparticle formation in the Pure Nano systems.

Building on these promising findings, more nanoparticles were prepared utilising different combinations of chemical molecules (molecular groupings, Table 3) with the aim of expanding the repertoire of accessible nanoparticles and further understanding the factors that influence their formation. Figure 1 in the Supplementary Information depicts characteristic dispersions and powders of several Pure Nano systems.

**Table 3:** Formulation, Size Distribution, and Surface Charge for Individual Pure Nano System.

| Molecular grouping | Molecule No. | [W]<sup>1</sup> | [R]<sup>2</sup> | [O]<sup>3</sup> | pH<sup>4</sup> | [W]<sup>5</sup> | Size<sub>(nm)</sub> | Zeta<sub>(mV)</sub> | PDI |
| --- | --- | --- | --- | --- | --- | --- | --- | --- | --- |
| 1 | 11, 6 | 3.0, 3.3 | 1:1 | 300 | 7 | 20 | 160.7 ± 1.785 | -48.5 ± 2.70 | 0.167 ± 0.047 |
| 2 | 11, 7 | 3.0, 2.7 | 1:1 | 300 | 7 | 20 | 139.3 ± 5.224 | -41.4 ± 2.48 | 0.132 ± 0.037 |
| 3 | 11, 8 | 3.0, 2.9 | 1:1 | 300 | 8 | 20 | 113.8 ± 3.978 | -32.9 ± 1.56 | 0.089 ± 0.097 |
| 4 | 11, 9 | 3.0, 4.9 | 1:1 | 300 | 9 | 20 | 157.3 ± 1.223 | -47.4 ± 1.47 | 0.162 ± 0.062 |
| 5 | 10, 9 | 3.0, 3.5 | 1:1 | 300 | 9 | 20 | 238.8 ± 4.809 | -54.6 ± 1.19 | 0.298 ± 0.005 |
| 6 | 10, 6 | 3.0, 2.4 | 1:1 | 300 | 7 | 20 | 166.3 ± 2.717 | -50.4 ± 3.34 | 0.177 ± 0.053 |
| 7 | 19, 10 | 3.0, 9.8 | 1:3 | 500 | 7 | 30 | 133.0 ± 7.169 | -39.3 ± 2.39 | 0.121 ± 0.081 |
| 8 | 20, 11 | 3.0, 7.5 | 1:5 | 500 | 9 | 35 | 120.9 ± 5.151 | -35.3 ± 3.20 | 0.101 ± 0.049 |
| 9 | 21, 11 | 3.0, 11.0 | 1:10 | 900 | 9 | 40 | 160.4 ± 1.749 | -48.4 ± 1.98 | 0.167 ± 0.051 |
| 10 | 22, 10 | 3.0, 3.8 | 1:3 | 300 | 7 | 20 | 151.6 ± 0.275 | -45.5 ± 1.50 | 0.152 ± 0.085 |
| 11 | 22, 11 | 3.0, 2.7 | 1:3 | 300 | 9 | 20 | 189.0 ± 6.512 | -58.0 ± 0.25 | 0.215 ± 0.072 |
| 12 | 23, 11 | 3.0, 4.0 | 1:3 | 300 | 9 | 20 | 128.2 ± 6.373 | -37.7 ± 1.47 | 0.113 ± 0.009 |
| 13 | 15, 12 | 3.0, 3.7 | 1:1 | 300 | 6 | 20 | 105.9 ± 2.658 | 30.3 ± 1.17 | 0.076 ± 0.039 |
| 14 | 15, 13 | 3.0, 3.4 | 1:1 | 300 | 6 | 20 | 97.80 ± 1.300 | 27.6 ± 2.00 | 0.063 ± 0.076 |
| 15 | 18, 12 | 3.0, 4.6 | 1:2 | 300 | 7 | 20 | 187.3 ± 6.220 | 57.4 ± 0.41 | 0.212 ± 0.052 |
| 16 | 18, 13 | 3.0, 4.2 | 1:2 | 300 | 7 | 20 | 170.1 ± 3.354 | 51.7 ± 1.80 | 0.183 ± 0.059 |
| 17 | 17, 12 | 3.0, 4.0 | 1:2 | 300 | 7 | 20 | 204.0 ± 9.005 | 63.0 ± 0.11 | 0.240 ± 0.020 |
| 18 | 17, 13 | 3.0, 3.7 | 1:2 | 300 | 7 | 20 | 114.0 ± 4.005 | 43.0 ± 0.11 | 0.090 ± 0.052 |
| 19 | 18, 10 | 3.0, 3.3 | 1:1 | 300 | 7 | 20 | 130.1 ± 6.690 | 38.3 ± 2.80 | 0.116 ± 0.054 |
| 20 | 18, 11 | 3.0, 2.8 | 1:1 | 300 | 7 | 20 | 80.20 ± 8.375 | 21.7 ± 1.50 | 0.033 ± 0.034 |
| 21 | 17, 8 | 3.0, 3.5 | 1:2 | 300 | 3 | 20 | 133.7 ± 7.299 | 46.5 ± 3.98 | 0.122 ± 0.026 |
| 22 | 17, 10 | 3.0, 5.0 | 1:2 | 400 | 6 | 25 | 158.1 ± 1.365 | 47.7 ± 2.31 | 0.163 ± 0.004 |
| 23 | 17, 11 | 3.0, 3.6 | 1:2 | 300 | 6 | 20 | 158.8 ± 1.469 | 47.9 ± 0.39 | 0.164 ± 0.062 |
| 24 | 19, 12 | 3.0, 2.6 | 1:1 | 300 | 7 | 20 | 142.1 ± 8.692 | -42.3 ± 0.85 | 0.136 ± 0.048 |
| 25 | 20, 13 | 3.0, 7.7 | 1:5 | 400 | 7 | 20 | 161.6 ± 1.944 | -48.8 ± 1.89 | 0.169 ± 0.083 |
| 26 | 21, 13 | 3.0, 11.2 | 1:10 | 700 | 7 | 30 | 169.0 ± 3.173 | -51.3 ± 0.46 | 0.181 ± 0.095 |
| 27 | 22, 13 | 3.0, 4.7 | 1:5 | 300 | 7 | 25 | 143.7 ± 7.288 | -49.5 ± 1.76 | 0.222 ± 0.054 |
| 28 | 22, 14 | 3.0, 3.6 | 1:5 | 300 | 8 | 25 | 95.10 ± 0.857 | -26.7 ± 0.15 | 0.058 ± 0.025 |
| 29 | 23, 14 | 3.0, 3.2 | 1:3 | 300 | 8 | 20 | 108.9 ± 3.163 | -31.3 ± 3.26 | 0.081 ± 0.054 |
| 30 | 12, 6 | 3.0, 2.9 | 1:1 | 300 | 7 | 20 | 129.5 ± 6.585 | -38.1 ± 2.70 | 0.115 ± 0.092 |
| 31 | 12, 7 | 3.0, 2.4 | 1:1 | 300 | 7 | 20 | 133.3 ± 7.219 | -39.4 ± 1.38 | 0.122 ± 0.048 |
| 32 | 13, 6 | 3.0, 3.2 | 1:1 | 300 | 7 | 20 | 153.7 ± 0.625 | -46.2 ± 1.51 | 0.156 ± 0.014 |
| 33 | 13, 7 | 3.0, 2.7 | 1:1 | 300 | 7 | 20 | 140.0 ± 8.336 | -41.6 ± 0.73 | 0.133 ± 0.067 |
| 34 | 14, 10 | 3.0, 5.3 | 1:1 | 300 | 3 | 20 | 181.3 ± 5.227 | 55.4 ± 1.55 | 0.202 ± 0.078 |
| 35 | 14, 11 | 3.0, 3.8 | 1:1 | 300 | 3 | 20 | 137.7 ± 7.957 | 40.9 ± 4.14 | 0.129 ± 0.007 |
| 36 | 15, 10 | 3.0, 4.6 | 1:1 | 300 | 6 | 20 | 157.1 ± 1.198 | 47.3 ± 2.97 | 0.161 ± 0.085 |
| 37 | 15, 11 | 3.0, 3.3 | 1:1 | 300 | 6 | 20 | 151.4 ± 0.248 | 45.4 ± 0.96 | 0.152 ± 0.083 |
| 38 | 16, 10 | 3.0, 4.9 | 1:1 | 300 | 7 | 20 | 158.8 ± 1.480 | 47.9 ± 1.61 | 0.164 ± 0.064 |
| 39 | 16, 11 | 3.0, 3.5 | 1:1 | 300 | 7 | 20 | 143.7 ± 8.954 | 42.9 ± 1.80 | 0.139 ± 0.044 |
| 40 | 24, 10 | 3.0, 4.1 | 1:1 | 300 | 7 | 20 | 139.6 ± 8.273 | 41.5 ± 3.47 | 0.132 ± 0.039 |
| 41 | 24, 11 | 3.0, 2.9 | 1:1 | 300 | 7 | 20 | 134.0 ± 7.348 | 39.6 ± 1.96 | 0.123 ± 0.084 |
| 42 | 26, 8 | 3.0, 2.4 | 1:1 | 300 | 3 | 20 | 145.8 ± 7.635 | 50.2 ± 2.71 | 0.226 ± 0.073 |
| 43 | 26, 11 | 3.0, 2.5 | 1:1 | 300 | 7 | 20 | 142.4 ± 8.738 | 42.4 ± 2.76 | 0.137 ± 0.066 |
| 44 | 24, 12 | 3.0, 3.3 | 1:1 | 300 | 7 | 20 | 168.2 ± 3.035 | 51.0 ± 0.70 | 0.180 ± 0.089 |
| 45 | 24, 13 | 3.0, 3.0 | 1:1 | 300 | 7 | 20 | 149.7 ± 9.951 | 44.9 ± 2.30 | 0.149 ± 0.096 |

Notes:

| Molecular grouping | Molecule No. | [W]<sup>1</sup> | [R]<sup>2</sup> | [O]<sup>3</sup> | pH<sup>4</sup> | [W]<sup>5</sup> | Size<sub>(nm)</sub> | Zeta<sub>(mV)</sub> | PDI |
| --- | --- | --- | --- | --- | --- | --- | --- | --- | --- |
| 46 | 26, 12 | 3.0, 2.8 | 1:1 | 300 | 7 | 20 | 157.1 ± 1.196 | 47.3 ± 3.93 | 0.161 ± 0.020 |
| 47 | 26, 13 | 3.0, 2.6 | 1:1 | 300 | 7 | 20 | 154.4 ± 0.747 | 46.4 ± 2.95 | 0.157 ± 0.000 |
| 48 | 24, 10, 12 | 3.0, 4.1, 3.3 | 1:1:1 | 500 | 7 | 25 | 93.60 ± 0.603 | 26.2 ± 1.70 | 0.056 ± 0.011 |
| 49 | 24, 10, 13 | 3.0, 4.1, 3.0 | 1:1:1 | 500 | 7 | 25 | 125.1 ± 5.860 | 36.7 ± 2.20 | 0.108 ± 0.095 |
| 50 | 24, 11, 12 | 3.0, 2.9, 3.3 | 1:1:1 | 500 | 7 | 25 | 127.2 ± 6.213 | 37.4 ± 3.27 | 0.112 ± 0.014 |
| 51 | 24, 11, 13 | 3.0, 2.9, 3.0 | 1:1:1 | 500 | 7 | 25 | 125.1 ± 5.852 | 36.7 ± 2.40 | 0.108 ± 0.058 |
| 52 | 26, 10, 13 | 3.0, 3.5, 2.6 | 1:1:1 | 500 | 7 | 25 | 149.5 ± 9.932 | 44.8 ± 3.64 | 0.149 ± 0.003 |
| 53 | 26, 11, 13 | 3.0, 2.5, 2.6 | 1:1:1 | 500 | 7 | 25 | 200.0 ± 8.349 | 61.6 ± 1.99 | 0.233 ± 0.059 |
| 54 | 1, 8 | 3.0, 4.9 | 1:1 | 300 | 9 | 20 | 134.2 ± 7.377 | -39.7 ± 4.55 | 0.123 ± 0.026 |
| 55 | 2, 6 | 3.0, 5.1 | 1:1 | 300 | 7 | 20 | 142.6 ± 8.775 | -42.5 ± 3.51 | 0.137 ± 0.014 |
| 56 | 2, 7 | 3.0, 4.2 | 1:1 | 300 | 7 | 20 | 158.4 ± 1.408 | -47.8 ± 1.17 | 0.164 ± 0.003 |
| 57 | 3, 6 | 3.0, 4.1 | 1:1 | 300 | 7 | 20 | 155.0 ± 0.839 | -46.6 ± 1.78 | 0.158 ± 0.026 |
| 58 | 4, 7 | 3.0, 3.1 | 1:1 | 300 | 7 | 20 | 142.2 ± 8.714 | -42.4 ± 1.29 | 0.137 ± 0.003 |
| 59 | 5, 6 | 3.0, 3.5 | 1:2 | 300 | 7 | 20 | 159.2 ± 1.539 | -48.0 ± 2.78 | 0.165 ± 0.091 |
| 60 | 1, 20 | 3.0, 2.0 | 5:1 | 300 | 7 | 20 | 85.10 ± 9.198 | -23.3 ± 1.97 | 0.041 ± 0.063 |
| 61 | 2, 20 | 3.0, 1.8 | 5:1 | 300 | 7 | 20 | 135.7 ± 7.620 | -40.2 ± 2.40 | 0.126 ± 0.082 |
| 62 | 2, 22 | 3.0, 3.0 | 5:1 | 300 | 7 | 20 | 140.2 ± 8.368 | -41.7 ± 1.37 | 0.133 ± 0.099 |
| 63 | 3, 19 | 3.0, 1.6 | 3:1 | 300 | 7 | 20 | 144.9 ± 9.154 | -43.3 ± 0.90 | 0.141 ± 0.096 |
| 64 | 4, 20 | 3.0, 3.4 | 2:1 | 300 | 7 | 20 | 138.7 ± 8.130 | -41.2 ± 1.60 | 0.131 ± 0.071 |
| 65 | 5, 23 | 3.0, 1.8 | 2:1 | 300 | 7 | 20 | 182.2 ± 5.374 | -55.7 ± 1.48 | 0.203 ± 0.068 |
| 66 | 1, 15 | 3.0, 4.6 | 1:1 | 300 | 6 | 20 | 177.2 ± 4.544 | 54.0 ± 3.88 | 0.195 ± 0.032 |
| 67 | 2, 16 | 3.0, 3.9 | 1:1 | 300 | 7 | 20 | 151.0 ± 0.171 | 45.3 ± 3.42 | 0.151 ± 0.069 |
| 68 | 3, 15 | 3.0, 3.4 | 1:1 | 300 | 6 | 20 | 148.3 ± 9.723 | 44.4 ± 2.47 | 0.147 ± 0.047 |
| 69 | 4, 16 | 3.0, 2.9 | 1:1 | 300 | 7 | 20 | 237.2 ± 4.541 | 74.0 ± 0.83 | 0.295 ± 0.063 |
| 70 | 5, 15 | 3.0, 1.4 | 1:1 | 300 | 6 | 20 | 138.2 ± 8.045 | 41.0 ± 2.90 | 0.130 ± 0.067 |
| 71 | 5, 16 | 3.0, 1.4 | 1:1 | 300 | 7 | 20 | 172.5 ± 3.761 | 52.5 ± 2.22 | 0.187 ± 0.065 |
| 72 | 1, 17 | 3.0, 1.7 | 5:1 | 300 | 6 | 20 | 149.4 ± 9.904 | 44.8 ± 2.90 | 0.149 ± 0.006 |
| 73 | 2, 17 | 3.0, 1.5 | 5:1 | 300 | 6 | 20 | 116.3 ± 2.226 | 49.4 ± 1.53 | 0.072 ± 0.097 |
| 74 | 2, 18 | 3.0, 1.4 | 5:1 | 300 | 7 | 20 | 116.5 ± 4.422 | 33.8 ± 0.44 | 0.094 ± 0.022 |
| 75 | 3, 17 | 3.0, 2.1 | 3:1 | 300 | 6 | 20 | 139.2 ± 8.214 | 41.4 ± 1.29 | 0.132 ± 0.054 |
| 76 | 4, 18 | 3.0, 2.5 | 2:1 | 300 | 7 | 20 | 118.0 ± 4.672 | 34.3 ± 2.45 | 0.096 ± 0.096 |
| 77 | 5, 18 | 3.0, 1.2 | 2:1 | 300 | 7 | 20 | 129.5 ± 6.594 | 38.1 ± 0.88 | 0.115 ± 0.046 |
| 78 | 1, 25 | 3.0, 1.5 | 5:1 | 300 | 7 | 20 | 173.2 ± 3.881 | 52.7 ± 2.63 | 0.188 ± 0.032 |
| 79 | 2, 24 | 3.0, 0.9 | 5:1 | 300 | 7 | 20 | 143.8 ± 8.975 | 42.9 ± 3.50 | 0.139 ± 0.076 |
| 80 | 2, 25 | 3.0, 1.4 | 5:1 | 300 | 7 | 20 | 141.0 ± 8.516 | 42.0 ± 1.32 | 0.135 ± 0.089 |
| 81 | 2, 26 | 3.0, 1.1 | 5:1 | 300 | 7 | 20 | 126.7 ± 6.121 | 45.9 ± 1.43 | 0.111 ± 0.088 |
| 82 | 4, 25 | 3.0, 2.6 | 2:1 | 300 | 7 | 20 | 137.2 ± 7.870 | 40.7 ± 1.41 | 0.128 ± 0.098 |

Notes:
Molecule No. in column 2 of each group correlates with the labelling system depicted in Table 1.
<sup>1</sup> Mass of the corresponding molecule (mg);
<sup>2</sup> Molar ratio of the molecules;
<sup>3</sup> Volume of organic solvent (μL)
<sup>4</sup> Acidity of aqueous solution. Buffer A, B, C, and D corresponded to sodium acetate buffer, cacodylate buffer, citrate buffer, and phosphate-citrate buffer, respectively;
<sup>5</sup> Volume of aqueous solution (mL).

### 3.3 Pure Nano Characterisation

#### 3.3.1 Size, Distribution, Zeta Potential, and Morphology

Figure 2 illustrates the representative size distributions, surface charges, and morphologies of nanoparticles formulated using different combinations of chemical molecules (molecular groupings), analysed through dynamic light scattering (DLS) and scanning electron microscope (SEM) techniques.

**Figure 2:**
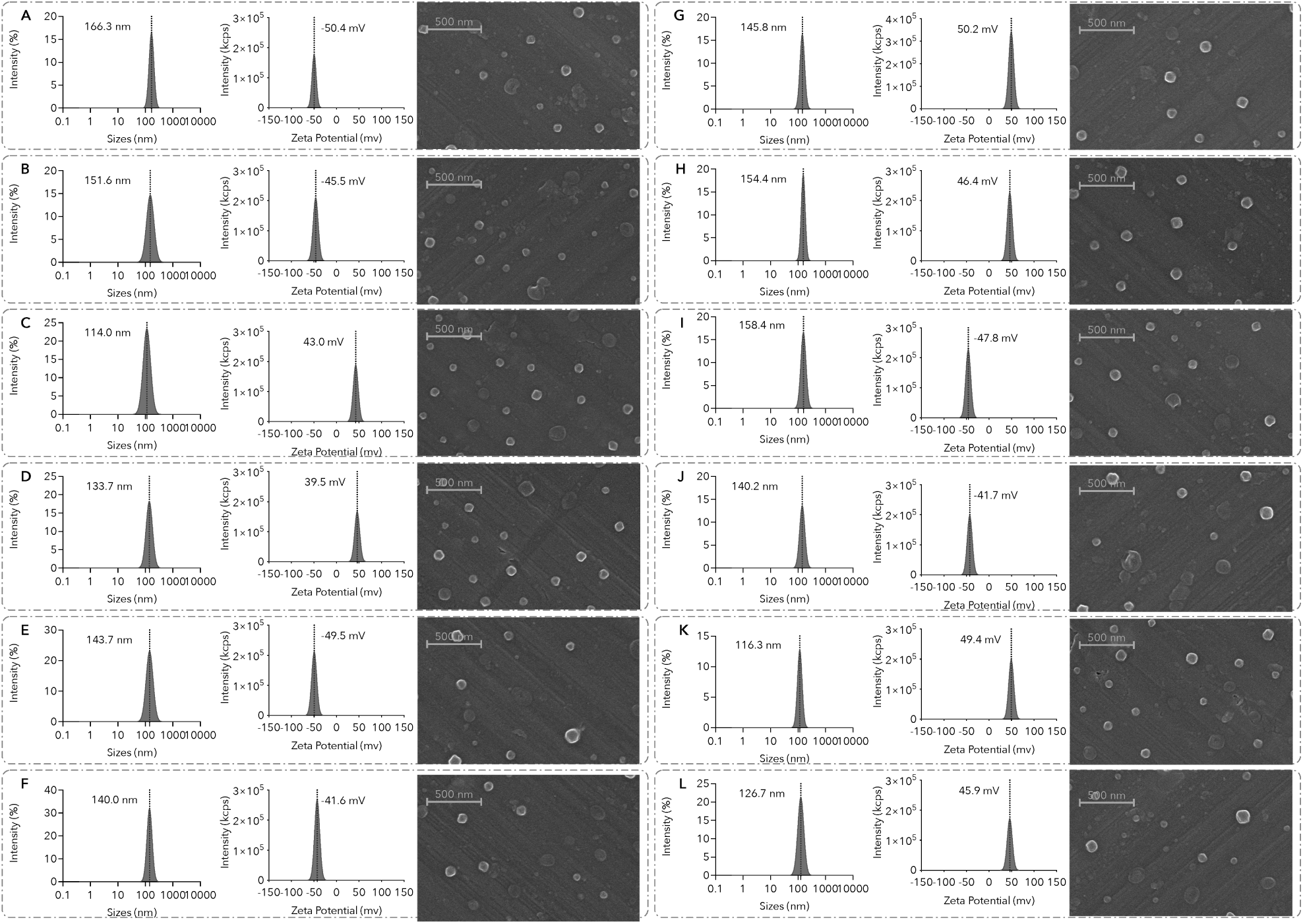
The representative size distribution, surface charges, and morphologies of Pure Nano systems formulated with (A) Naphthofluorescein and aristolochic acid, grouping number 6 in Table 3, (B) Rose bengal sodium and naphthofluorescein, grouping number 10 in Table 3, (C) Mitoxantrone dihydrochloride and nile red, grouping number 18 in Table 3, (D) Mitoxantrone dihydrochloride and ellagic acid, grouping number 21 in Table 3, (E) Rose bengal sodium and nile red, grouping number 27 in Table 3, (F) Rhein and nile red, grouping number 33 in Table 3, (G) Berberine chloride and ellagic acid, grouping number 42 in Table 3, (H) Berberine chloride and nile red, grouping number 47 in Table 3, (I) Pyrene and rhein, grouping number 56 in Table 3, (J) Pyrene and rose bengal sodium, grouping number 62 in Table 3, (K) Mitoxantrone dihydrochloride and pyrene, grouping number 73 in Table 3, (L) Berberine chloride and pyrene, grouping number 81 in Table 3.

These molecular groupings include the combination of: (A) weak acids (Naphthofluorescein and aristolochic acid, grouping number 6 in Table 3); (B) weak acids and conjugate bases of weak acids (Rose bengal sodium and naphthofluorescein, grouping number 10 in Table 3); (C) weak bases and conjugated acids of weak bases (Mitoxantrone dihydrochloride and nile red, grouping number 18 in Table 3); (D) weak acids and conjugated acids of weak bases (Mitoxantrone dihydrochloride and ellagic acid, grouping number 21 in Table 3); (E) weak bases and conjugated bases of weak acids (Rose bengal sodium and nile red, grouping number 27 in Table 3); (F) weak acids and weak bases (Rhein and nile red, grouping number 33 in Table 3; (G) permatently charged molecules and weak acids (Berberine chloride and ellagic acid, grouping number 42 in Table 3); (H) permatently charged molecules and weak bases (Berberine chloride and nile red, grouping number 47 in Table 3); (I) non-ionisable molecules and weak acids (Pyrene and rhein, grouping number 56 in Table 3); CT) non-ionisable molecules and conjugated bases of weak acids (Pyrene and rose bengal sodium, grouping number 62 in Table 3); (K) non-ionisable molecules and conjugated acids of weak bases (Mitoxantrone dihydrochloride and pyrene, grouping number 73 in Table 3); (L) permatently charged molecules and non-ionisable molecules (Berberine chloride and pyrene, grouping number 81 in Table 3).

As can be seen, the Pure Nano systems exhibit hydrodynamic sizes between 100 nm to 150 nm with narrow distributions, alongside high absolute surface charge values exceeding 40 mV. These attributes are essential for their formation and potential dispersion stability as a high zeta potential (typically ±30 *mV* or higher) is an indication of good dispersion and stability of nanoparticles in a colloidal dispersion.^44^ Besides, SEM analyses confirmed uniform dispersion and revealed regular spherical shapes of the nanoparticles within the desired nanometre range.

#### 3.3.2 Fluorescence

The excitation, emission and synchronous fluorescence spectra of PyRhe Nano (formulated with pyrene and rhein) by re-dispersing them into water with varying quantities of DMSO were displayed in Figure 3A, B and C, respectively. As can be seen, the excitation spectra reveal multiple absorption peaks, ranging from 260 nm to 400 nm, with a peak around 395 nm displaying increased intensity upon reducing the concentration of DMSO in the solvent. Simultaneously, other peaks show decreased intensity, indicating a red-shift in the absorption spectra of the nanoparticles with decreasing DMSO concentration. The emission spectra reveal not only visible-range fluorescence but also fluorescence beyond 900 nm, with intensities up to 100 times higher than those in the visible range, implying substantial changes in the fluorescence properties caused by the nanoparticle formation.

**Figure 3:**
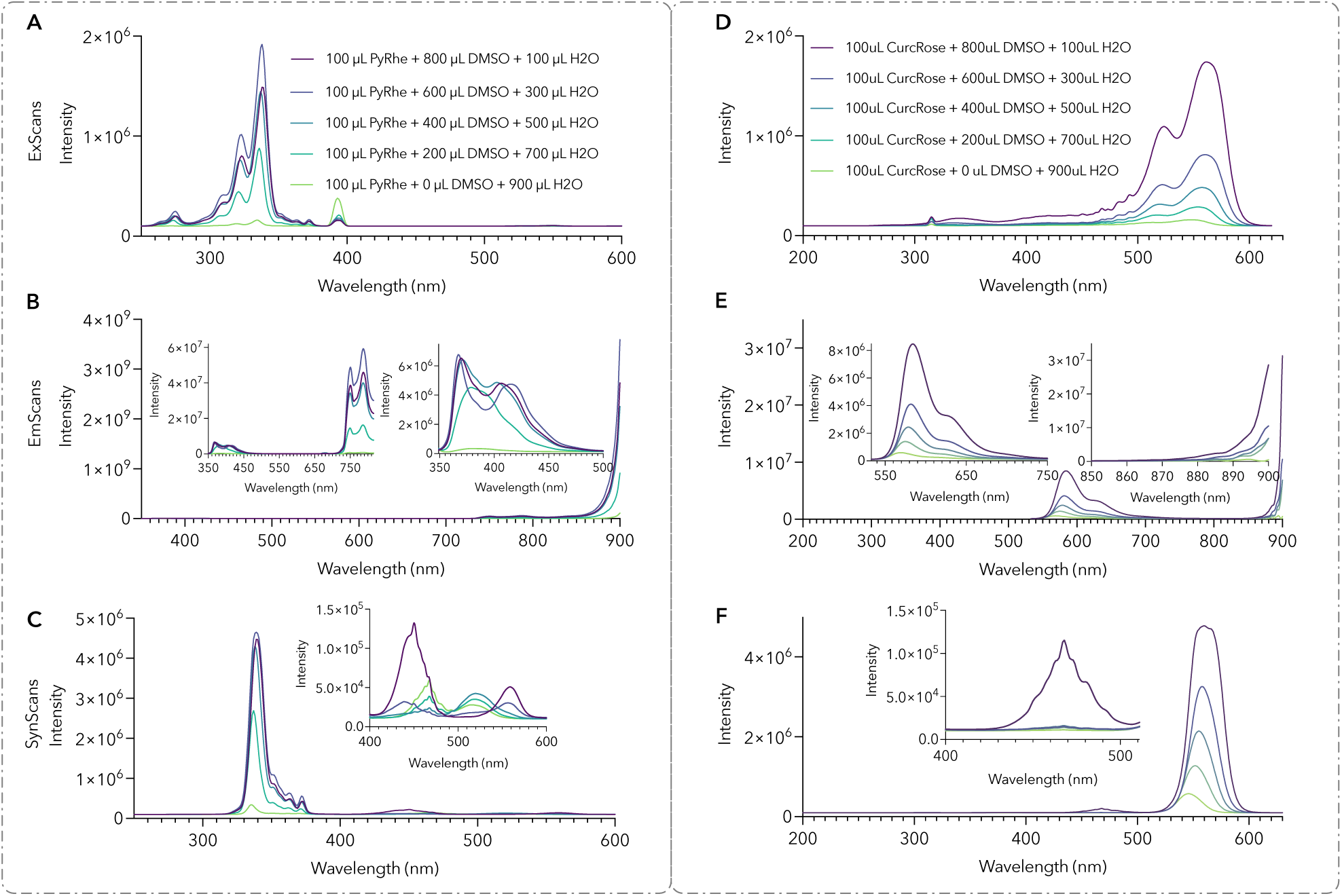
(A) PyRhe Nano (formulated with pyrene and rhein) displays multiple absorption peaks ranging from 260 nm to 400 nm in their excitation spectra. The peak intensity at 395 nm increases upon reducing the concentration of DMSO while other peaks show decreased intensity. Their emission spectra (B) reveal both visible and NIR fluorescence while synchronous fluorescence spectra (C) shows multiple fluorescent species in the nanoparticles. (D) CurcRose Nano (formulated with curcumin and rose bengal) exhibits significant visible absorption, ranging from around 500 nm to about 600 nm, with a peak at 545 nm. And they emit fluorescent radiation in both visible range (550 nm to 700 nm) and NIR range (E). The synchronous fluorescence spectra (F) of CurcRose Nano dispersed in water exhibit only one emission peak at 545 nm, suggesting the presence of a single fluorescent species. The variation in DMSO concentration leads to dramatical changes in both the absorption and emission intensities of the nanoparticles.

The synchronous fluorescence spectrum is measured by concurrently scanning the excitation and emission wavelengths while using the peak wavelength difference between the spectra, which has been extensively used in multicomponent analysis to discriminate between different fluorescent species based on their distinctive fluorescence properties.^45^ The synchronous fluorescence spectra of each fluorescent species in the mixture will have a distinct peak shape and position, allowing for identification and quantification of each component. As depicted in Figure 3C, the nanoparticles in water exhibit three distinct peaks, centred around 350 nm, 470 nm, and 520 nm, respectively. The changes of fluorescent intensities and shifts in peak positions, as induced by the concentration variation of DMSO in the solvent, could potentially result from the differences in strength and degree of aggregation during nanoparticle formation under different conditions, which produced different fluorescent species.

Similarly, rose bengal pyrene nanoparticles exhibit a nearly identical excitation and emission behaviour. As shown in Figure 3D, the nanoparticles in water exhibit significant visible absorption, ranging from around 500 nm to about 600 nm, with a peak at 545 nm. As the concentration of DMSO in the solvent increases, the absorption intensity and width increase, and the 550 nm peak splits into two peaks at around 525 nm and 565 nm. The nanoparticles emit fluorescent radiation in both visible range (550 nm to 700 nm) and NIR range (beyond 900 nm), as shown in Figure 3E. The synchronous fluorescence spectra of rose bengal pyrene nanoparticles dispersed in water exhibit only one emission peak at 545 nm (Figure 3F), suggesting the presence of a single fluorescent species. However, introducing a sufficient amount of DMSO into the solvent may modify the size or morphology of the nanoparticles, resulting in their inhomogeneity and multiple peaks with a red-shift.

#### 3.3.3 Stability

To further verify the stability of various Pure Nano systems, such as PyTop (formulated with pyrene and topotecan hydrochloride, Figure 4A) in water, PyRhe (formulated with pyrene and rhein, Figure 4B) in PBS (*pH*, 7.4) and PyMB (formulated with pyrene and methylene blue, Figure 4B) in water, their changes in sizes, distributions, and surface charges were measured under different conditions.

**Figure 4:**
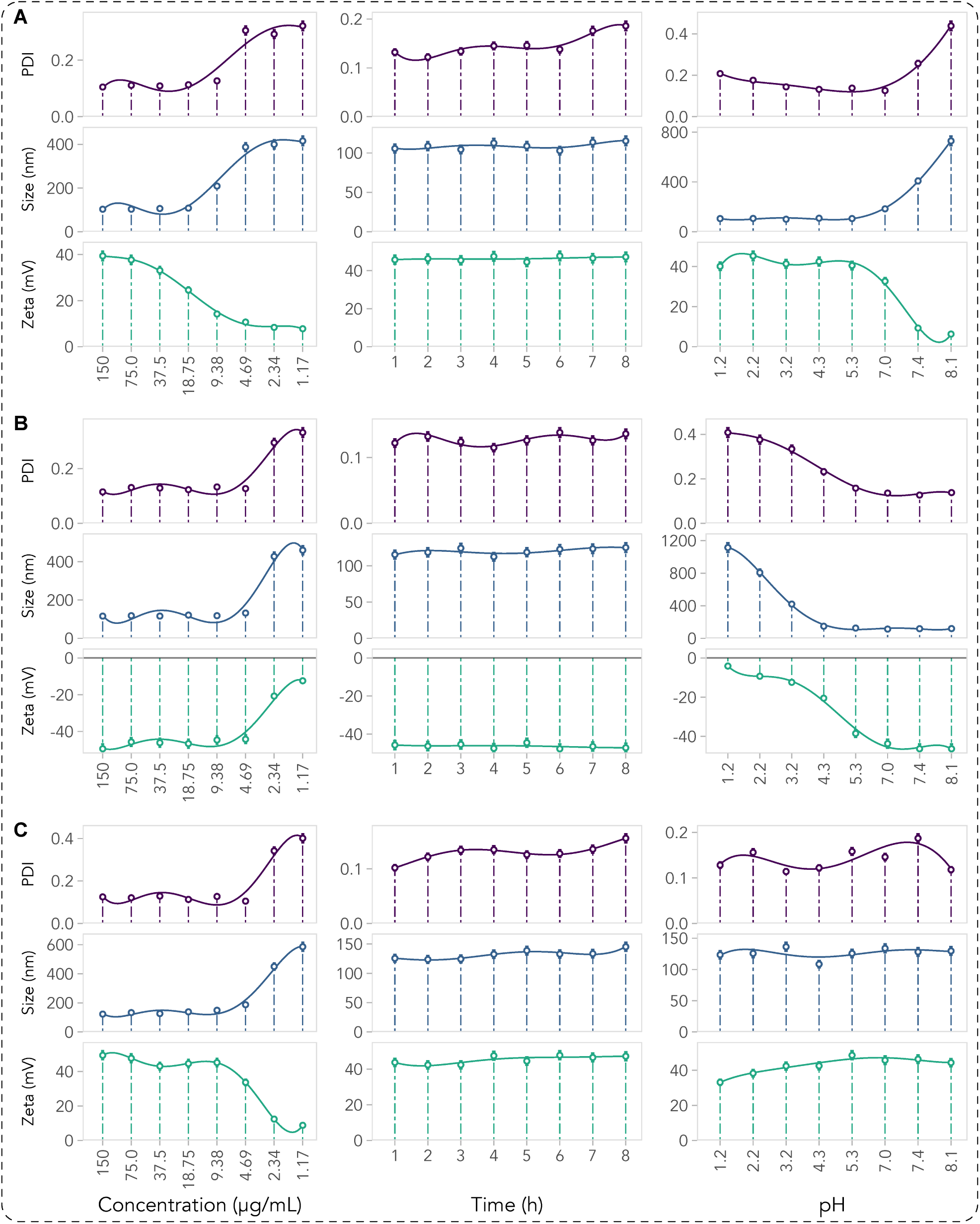
The sizes, distributions (PDI), and surface charges of various Pure Nano systems including (A) PyTop (formulated with pyrene and topotecan hydrochloride) in water (B) PyRhe (formulated with pyrene and rhein) in PBS *(pH,* 7.4, 150 µg/mL) (C) PyMB (formulated with pyrene and methylene blue, 150 µg/mL) in water can be sustained at a low concentration and maintained under preparative conditions in a 4°C environment for 8 hours. When the acidity of the aqueous solutions was manipulated, the particle size, distribution and surface charge of PyTop and PyRhe underwent significant changes, while PyMB remained stable.

**Figure 5:**
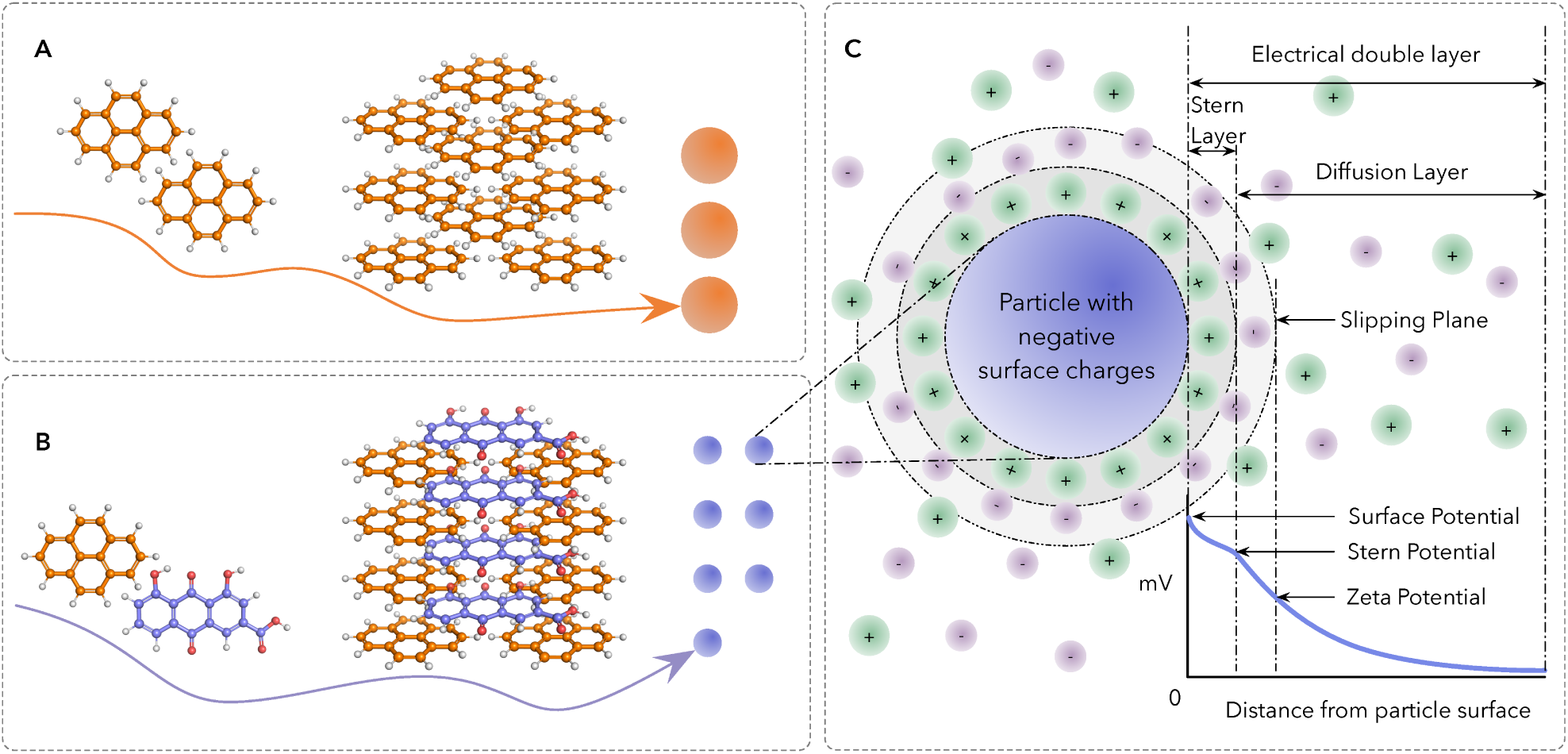
(A) An illustration depicting the spontaneous aggregation of conjugated molecules into particles. This aggregation process is driven by the strong intermolecular forces resulting from the π - π stacking interactions between the conjugated systems when they come into con­ tact with each other, leading to a decrease in free energy and an increase in size. (B) Spontaneous aggregation of specified conjugated molecules into nanoparticles with strong surface charges, providing repulsive forces to prevent further aggregation during the particle formation pro­ cess and leading to a uniform dispersion of particles with desired nanometre sizes. (C) An illustration of electrical double layer explaining the interaction of ions and charged particles at the interface of a liquid surface with an electrolyte solution.

As depicted, the sizes, distributions, and surface charges of nanoparticles could be sustained at a low concentration by reducing their dispersion concentration, indicating their excellent resistance to dilution. Moreover, the anti-dilution abilities of the nanoparticle systems with different components varied due to the differences in the intermolecular strength within different systems. The nanoparticle dispersions were maintained under preparative conditions in a 4°C environment for 8 hours, during which there were no evident changes detected in particle size, distribution, and surface charges, highlighting their excellent stability under the specified conditions.

However, upon manipulating the acidity of the aqueous solutions, the particle size, distribution, and surface charges of PyTop and PyRhe underwent significant changes at certain critical points. When the acidity exceeded these critical points, the nanoparticles experienced increased sizes, wider distributions, and a considerable decrease in surface charge from approximately ±50 *mv* to a magnitude of just a few millivolts. These critical points corresponded directly to the *pK_a_* values of the constituent molecules for the construction of the nanoparticle systems, thus emphasising the significance of surface charges in the establishment and stability of the nanoparticle systems.

Moreover, the methylene blue pyrene nanoparticles displayed excellent resistance to environmental disruptions due to the inclusion of both methylene blue and pyrene. Methylene blue is a permanently charged molecule that contains a positively charged nitrogen atom whose charge is dispersed throughout the entire structure, and pyrene is a non-ionisable molecule. The charge delocalisation of methylene blue and the non-ionisable property of pyrene endow the methylene blue pyrene nanoparticles with the ability to sustain their sizes and distributions without any disturbance from changes in the external environment, such as the environmental *pH* values (Figure 4C). Thus, remarkably high surface charges significantly contribute to the formation and stability of Pure Nano systems. Achieving this relies on selecting molecules with specific *pK_a_* values and modulating the corresponding acidity of aqueous solutions.

### 3.4 Formation Mechanism: Charge Balanced Aggregation of Conjugated Molecules

Zeta potential (() denotes the electric charge surrounding colloidal particles dispersed in a suspension or dispersion. **It** quantifies the potential difference between the stationary layer of fluid linked to the particle surface and the dispersion medium^46^, as shown in Figure SC.

The cause underlying zeta potential in particles lies in the presence of charged surface groups, which interact with electrolyte ions in the surrounding medium, leading to the formation of an electrical double layer around the particle. The double layer of charged particles surrounding the particle surface creates an electric potential difference between the stationary layer of fluid attached to the particle surface and the surrounding dispersion medium, known as the zeta potential.

In the process of spontaneous aggregation of conjugated molecules, the dissociation of ionisable conjugated molecules under environments with certain *pH* values provides the sources of charged groups, resulting in the development of a zeta potential on initially formed particles due to aggregation, as shown in figure SB. This also provides repulsive forces to prevent further aggregation of conjugated molecules. When the spontaneous aggregation is balanced by the repulsive forces provided by the surface charges, namely, charge balanced aggregation, a uniform dispersion of particles with desired sizes is attained.

Therefore, dissolving conjugated molecules with distinct dissociation types and degrees into aqueous environments at specific pH levels drives the formation of Pure Nano systems. Basically, an optimal quantity of ionised conjugated molecules is required to generate sufficiently robust repulsive forces that counterbalance the spontaneous aggregation of conjugated molecules in aqueous settings. To this end, a detailed framework for categorising the aggregation of conjugated molecules with different dissociation types and degrees is provided, as shown in Figure 6 from panel B to panel O.

**Figure 6:**
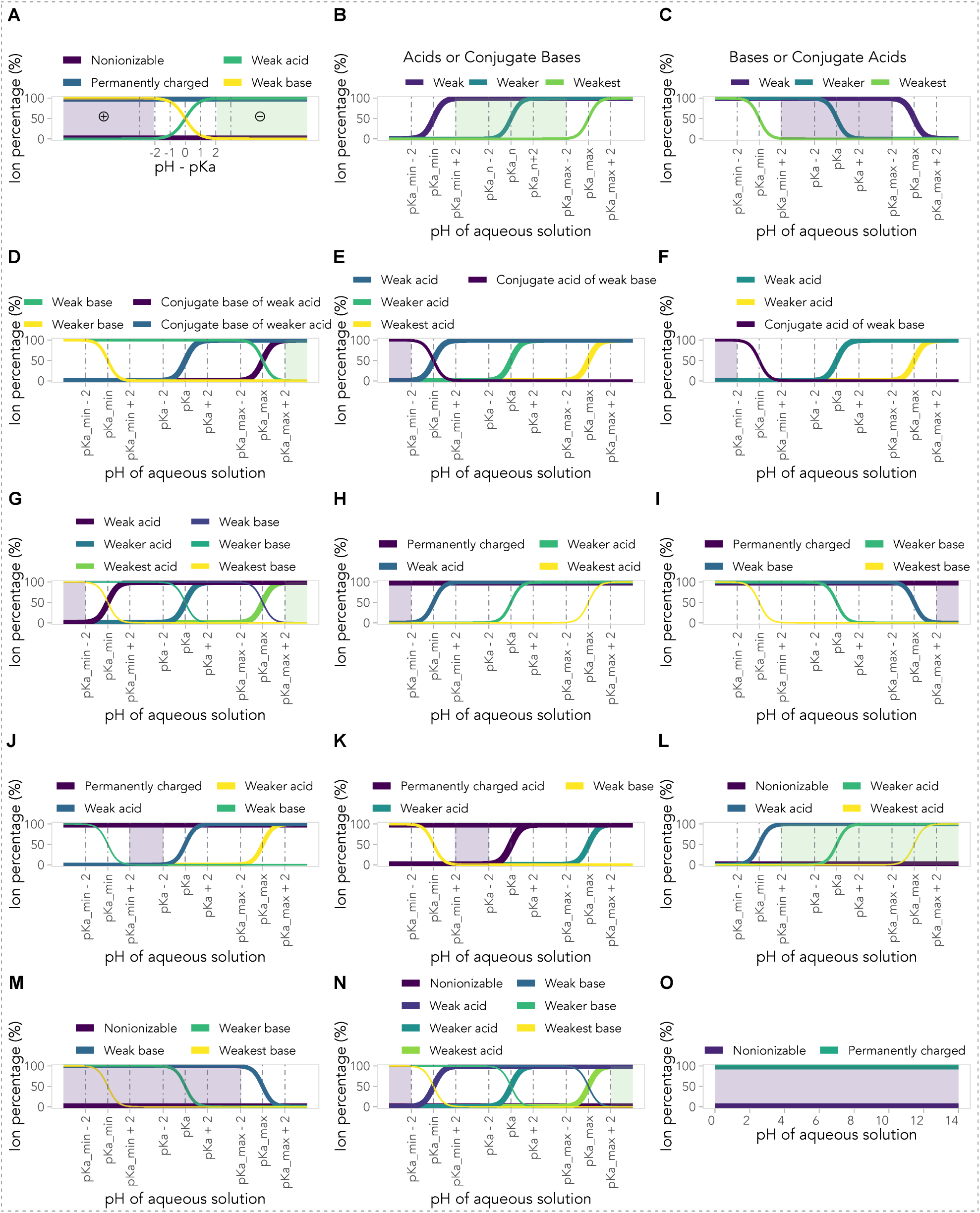
(A) The dissociation of molecules in buffering solutions at equilibrium. In an aqueous solution with a *pH* two units above the *pK_a_* value of a weak acid, around 99% of molecules will exist in a negatively charged ionic state. Conversely, in a solution with a *pH* two units below the *pK_a_* value ofa weak base, over 99% of molecules will be in a positively charged ionic state. (B-O) The*pH* ranges of aqueous solutions (shaded areas), the *pK_a_* relationships and the types of conjugated molecules utilised for the construction of distinct Pure Nano systems.

#### 3.4.1 Weak acids or conjugate bases of weak acids

Figure 6B depicts the state of a Pure Nano system consisting of multiple weak acids or conjugate bases of weak acids within an aqueous solution. Among these acids or conjugate bases, the one with the lowest *pK_a_* value (*pK_a_*__*min*_) demonstrates comparatively stronger acidity than the others, while the acid with the highest *pK_a_* value (*pK_a_*__*max*_) is the weakest in terms of acidity. The weak acid showing stronger acidity has a greater tendency to donate a proton (*H*^+^) than weaker ones, resulting in an easier dissociation under the same condition. To ensure sufficient dissociation of the weak acid and maintain the majority of the weakest acid in its molecular state (as depicted in the shade of Figure 6B), the *pH* of the aqueous solution should meet certain requirements, as shown in Equations 5.

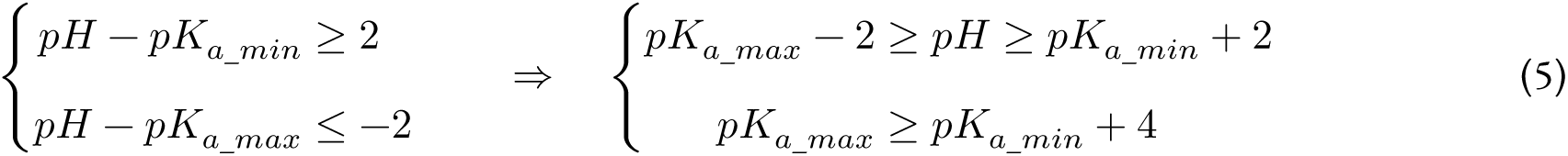

When the *pH* of the aqueous solution is two units larger than the *pK_a_* value of the weak acid (*pK_a_*__*min*_) , over 99% of the weak acid will dissociate into its ionic form. Conversely, if the *pH* of the solution is two units less than the *pK_a_* value of the weakest acid (*pK_a_*__*max*_), most of the weakest acid will remain in its molecular form. Thus, a molecular system with a *pH* value within the *pK_a_* range of the acids and at least two units away from the lowest and highest *pK_a_* values ensures the dissociation of the majority of the weak acid while maintaining the integrity of the weakest acid. And furthermore, to construct such a system, the *pK_a_* value of the weak acid (*pK_a_*__*min*_) should be at least 4 units lower than that of the weakest one (*pK_a_*__*max*_).

The highly negatively charged surface (high ζ values) of nanoparticles allows them to be uniformly (extremely small PDI values) dispersed in aqueous solutions, resulting in a stable colloidal dispersion. The pertinent nanoparticles acquired using the molecules from this grouping is listed in Table 3, with grouping numbers ranging from 1 to 12.

#### 3.4.2 Weak bases or conjugate acids of weak bases

Equations 5 are also suitable for systems comprising multiple weak bases or conjugate acids of weak bases in aqueous solutions. Aqueous solutions with a *pH* value within two units above the smallest *pK_a_* value of the weakest base (*pK_a_*__*min*_) and below the largest *pK_a_* value of weak base (*pK_a_*__*max*_) would allow for the dissociation of the majority of the weak bases while preserving the molecular state of the weakest base (as depicted in the shade of Figure 6C). Thus, the *pK_a_* value of the weak base (*pK_a_*__*max*_) should be at least 4 units above that of the weakest one (*pK_a_*__*min*_).

In such a Pure Nano system of conjugated basic molecules, the nanoparticles would be uniformly dispersed in aqueous solutions due to their highly positively charged surface. The relevant nanoparticles obtained by utilising molecules from this category are listed in Table 3 (grouping number from 13 and 18).

#### 3.4.3 Weak bases and conjugate bases of weak acids

For a system comprising weak bases and conjugate bases of weak acids (Figure 6D), Equation 6 illustrates the necessary conditions for the aqueous solution. To ensure that the conjugate bases of acids remain in ionic form in this system, the *pH* of the aqueous solution should be at least two units higher than their *pK_a_* values. Similarly, to maintain the weak bases in molecular form, the *pH* should also be at least two units higher than their *pK_a_* values. Therefore, the *pH* of the aqueous solution should be at least two units higher than the largest *pK_a_* value (*pK_a_*__*max*_) of the conjugated molecules to keep them in the desired forms in the system (Equations 6). In this case, there is no specific requirement of the *pK_a_* values between the conjugated molecules.

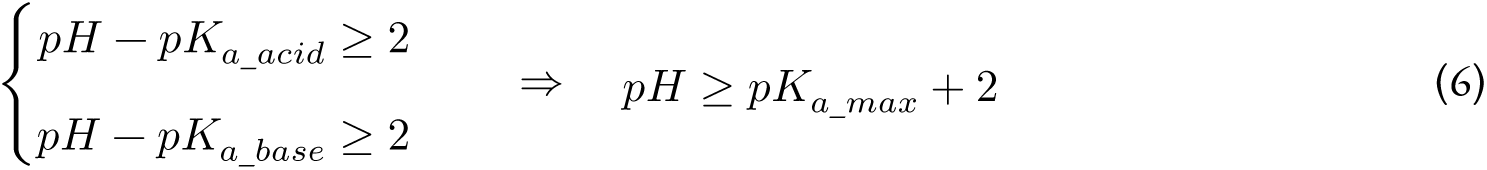

And in such a Pure Nano system, the surface of nanoparticles would be highly negatively charged, which ensures their uniform dispersion in solutions. Table 3 enumerates the relevant nanoparticles obtained using molecules from this grouping, with group numbers ranging from 24 to 29.

#### 3.4.4 Weak acids and conjugate acids of weak bases

Similarly, for a system containing weak acids and conjugate acids of weak bases (Figure 6E and F), the necessary conditions for the aqueous solution is shown in Equations 7. To keep the conjugate acids of weak bases in their ionic forms while maintain the integrity of the weak acids, the *pH* of the aqueous solution should be at least two units lower than the smallest *pK_a_* value (*pK_a_*__*min*_) of the conjugated molecules. And there is no specific requirement of the *pK_a_* values between the conjugated molecules. The formed nanoparticles would be highly positively charged, as listed in Table 3 with grouping numbers from 19 to 23.

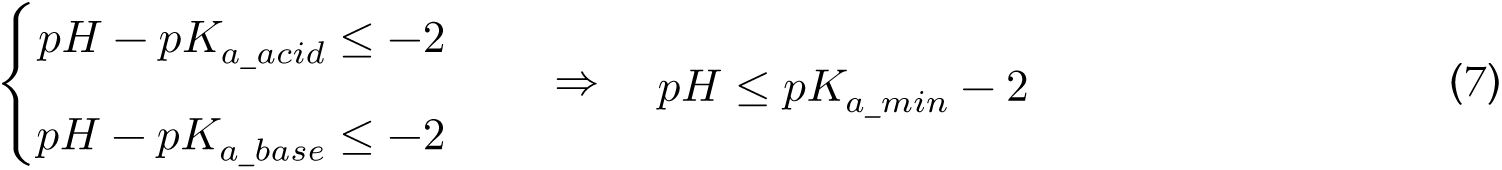

#### 3.4.5 Weak acids and weak bases

Figure 6G shows a system containing both weak acids and weak bases, which could form nanoparticles with positive charges or negative charges. To ensure the dissociation of weak bases and the integrity of weak acids, the *pH* of aqueous solutions should be at least two units lower than the smallest *pK_a_* value (*pK_a_*__*min*_) of the conjugated molecules (Equations 7), which allows the formation of positively charged nanoparticles in dispersion. Conversely, to allow the dissociation of weak acids and the integrity of weak bases, the *pH* of aqueous solutions should be at least two units higher than the largest *pK_a_* value (*pK_a_*__*max*_) of the conjugated molecules (Equations 6), resulting the negatively charged nanoparticles in dispersion. Table 3 presents the relevant nanoparticles obtained using molecules from this grouping, with group numbers ranging from 30 to 39.

#### 3.4.6 Permanently charged molecules and weak acids/bases

Permanently charged molecules are those that maintain their charges regardless of the *pH* of the surrounding environment. They are usually positively charged, such as methylene blue and berberine chloride used in this research. Thus, for systems that utilise permanently charged molecules as one of the primary components, it is only necessary to control the formation of the other molecules to achieve a Pure Nano system.

The formed nanoparticles will be highly positively charged in these systems.

For a system containing permanently charged molecules and weak acids, the *pH* of aqueous solutions should be at least two units lower than the smallest *pK_a_* value (*pK_a_*__*min*_) of the weak acids (Equations 8) to prevent their dissociation and maintain them in molecular forms (as depicted in the shade of Figure 6H). Otherwise, the dissociated acid would electro-statically interact with the permanently charged molecules, resulting in large particles in aqueous solutions. The nanoparticles obtained using molecules from this grouping are listed in Table 3, featuring group numbers ranging from 40 to 43.

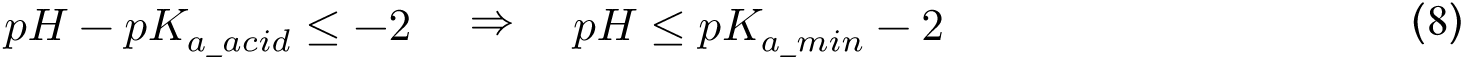

Conversely, for a system containing permanently charged molecules and weak bases (Figure 6I), the dissociation of weak bases should be controlled in order to achieve a charge balanced system. The *pH* of aqueous solutions should be at least two units larger than the largest *pK_a_* value (*pK_a_*__*max*_) of the weak bases (Equations 9). Table 3 presents the nanoparticles obtained through the utilisation of molecules from this grouping, with group numbers ranging from 44 to 47.

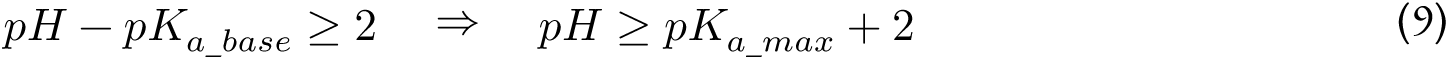

If permanently charged molecules are incorporated into a system that includes both weak acids and weak bases as the third dominant component (as depicted in Figure 6J), the dissociation of both acids and bases should be hindered by controlling their environments in order to achieve a Pure Nano system. The *pH* of aqueous solutions should be at least two units lower than the *pK_a_* value of the weak acids (*pK_a_*__*acid*_) while larger than the *pK_a_* value of the weak bases (*pK_a_*__*acid*_), as shown in Equations 10. Thus, the the *pK_a_* value of weak acids (*pK_a_*__*acid*_) should be at least 4 units above that of weak bases (*pK_a_*_*_base*_). The nanoparticles obtained through the usage of molecules from this grouping are listed in Table 3, featuring group numbers ranging from 48 to 53. Furthermore, if the permanently charged molecules contain ionisable acidic groups, they should also be regarded as acids (Figure 6K), and their *pK_a_* values should conform to Equations 10.

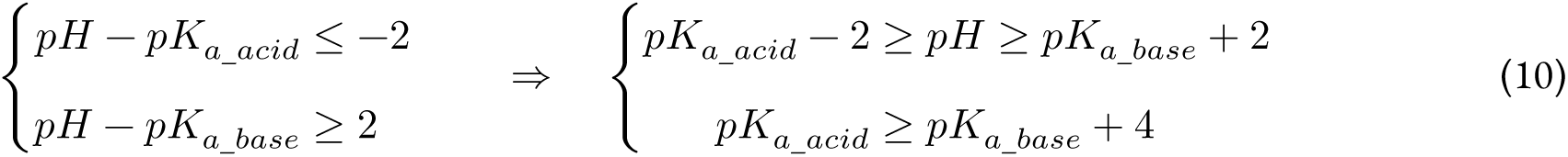

#### 3.4.7 Non-ionisable molecules and weak acids/bases

The introduction of non-ionisable molecules as one of the dominant components in a system requires only the control of the other conjugated molecules to provide the necessary charge for achieving a Pure Nano system. In a system utilising non-ionisable molecules and weak acids (depicted in Figure 6L), it is crucial to elevate the *pH* of aqueous solutions by at least two units above the minimum *pK_a_* value of the weak acids (*pK_a_*__*min*_) to ensure adequate formation of ionic acids (as described in Equations 11). The pertinent nanoparticles acquired using the molecules from this grouping is listed in Table 3, with grouping numbers ranging from 54 to 65.

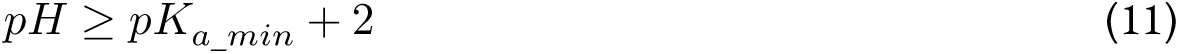

Conversely, in a system employing non-ionisable molecules and weak bases (as shown in Figure 6M), the *pH* of aqueous solutions must be lowered by at least two units below the maximum *pK_a_* value of the weak bases (*pK_a_*__*max*_) to achieve sufficient formation of ionic bases (as described in Equations 12).

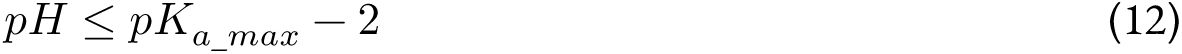

Furthermore, if non-ionisable molecules are introduced into a system predominantly comprising weak acids and weak bases (Figure 6N), the conditions (Equations 6 and 7) applicable to systems containing only weak acids and weak bases (Figure 6G) can also be applied to this system without any modifications. The pertinent nanoparticles acquired using the molecules from this grouping is listed in Table 3, with grouping numbers ranging from 66 to 77.

#### 3.4.8 Permanently charged molecules and non-ionisable molecules

The simplest Pure Nano system comprises permanently charged molecules and non-ionisable molecules (depicted in Figure 6O). The formation of aforementioned systems necessitates the control over the molecules and aqueous solution. However, in a system predominantly composed of permanently charged molecules and non-ionisable molecules, no such requirements exist, as their dissociation state remains unaffected by the environmental conditions or the *pK_a_* values of the molecules. The resulting nanoparticles would exhibit a high positive charge, irrespective of any fluctuations in the environmental *pH* values. The pertinent nanoparticles acquired using the molecules from this grouping is listed in Table 3, with grouping numbers ranging from 78 to 82.

### 3.5 Applications

#### 3.5.1 PyRhe Nano restored DSS-induced acute ulcerative colitis in mice

Ulcerative colitis, an inflammatory bowel disease primarily affecting the colon and rectum, constitutes a chronic condition typified by enduring inflammation and the development of ulcers along the gastrointestinal mucosa. Instances of abrupt and severe symptomatology in ulcerative colitis are identified as acute ulcerative colitis (UC), signifying an escalation in disease manifestation.

The efficacy of PyRhe Nano, formulated with pyrene and rhein, in mitigating UC induced by dextran sulfate sodium (DSS) was assessed in mice. As shown in Figure 7A, the colon length of UC mice was significantly shortened compared to the healthy control group. The free rhein molecule was found to significantly alleviate colon atrophy. Fascinatingly, the introduction of PyRhe Nano restored colon length to a level remarkably close to that of the healthy control group (no statistically significant difference found), demonstrating the pronounced protective effects against UC at the level of biological organs. The representative *ex vivo* mouse colon tissues that received different treatments are shown in Figure 2 in the Supplementary Information.

**Figure 7:**
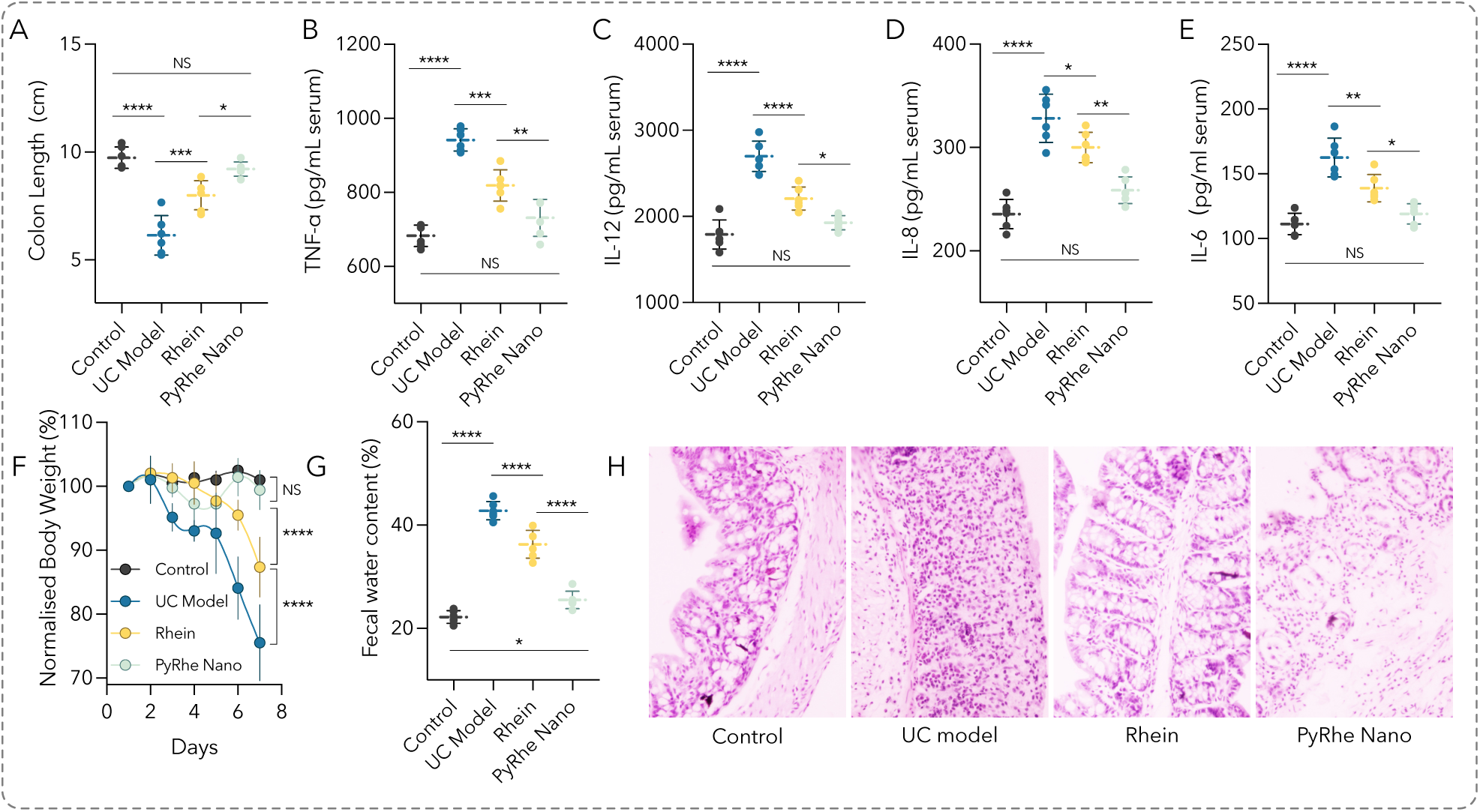
PyRhe Nano, formulated with pyrene and rhein, restored dextran sulfate sodium (DSS) induced acute ulcerative colitis in mice: reinstatement of colon length (A), normalisation of inflammatory factors including tumour necrosis factor-alpha (TNF-α), interleukin 6 (IL-6), interleukin 8 (IL-8) and interleukin 12 (IL-12), recovery from body weight loss (F) and water loss (G) comparable to healthy control group levels, demonstrating the significant protective effects. (H) Hematoxylin-eosin (H&E) staining displays tissue recovery in the PyRhe Nano group akin to the healthy control. Number of mice in each group: n = 6. * p < 0.05; ** p < 0.01; *** p < 0.001; **** p < 0.0001. H&E original magnification, 4X.

Similarly, the levels of inflammatory factors including TNF-*α* (Figure 7B), IL-12 (Figure 7C), IL-8 (Figure 7D), and IL-6 (Figure 7E) were significantly increased in response to DSS in UC mice. The administration of free rhein significantly decreased the levels of inflammatory factors, while PyRhe Nano restored the factors to the level of the healthy control group (no statistically significant difference found), showing a great protective effect of the nano formulation against UC at the molecular level.

Besides, the body weight of mice from model group and the group treated with free rhein decreased by approximately 25% and 15%, respectively (Figure 7F). Mice with UC that were treated with PyRhe Nano experienced a slight (5%) initial loss of body weight, but subsequently regained weight to the level of the healthy group, indicating a reversal of weight loss caused by UC. In addition, the water content of feces in UC mice increased significantly (Figure 7G), leading to severe diarrhea. The free rhein molecule also significantly increased fecal water content, revealing its disadvantages in diarrhea. However, administration of PyRhe Nano reduced fecal water content comparable to the healthy control group, suggesting a potential advantage in mitigating this aspect of diarrhea.

Figure 7H displays the H&E staining of the colon. In UC mice, DSS induced significant mucosal damage, characterized by crypt loss, focal necrosis, and infiltration of inflammatory cells into the colonic tissue. Notably, tissues from the free rhein molecule and PyRhe Nano groups exhibit H&E patterns similar to the healthy control group, serving as direct visualisation of the protective effects conferred by this nano formulation. All findings collectively indicate the PyRhe Nano’s effective protective role against DSS-induced acute ulcerative colitis in mice.

#### 3.5.2 CurcRhe Nano Reversed Myocardial Ischemia/Reperfusion (I/R) Injury in Mice

Ischemia/reperfusion (I/R) denotes the return of blood supply to tissue subsequent to a period of oxygen deprivation (ischemia). Despite being essential for tissue viability, the reinstatement of blood flow can paradoxically induce additional harm. The abrupt reintroduction of oxygen triggers an inflammatory cascade, accompanied by the generation of reactive oxygen species, culminating in tissue damage that exacerbates the initial injury.

As depicted in Figure 8A, extensive areas of white necrosis and cardiomyocyte edema were observed in I/R heart sections^47^ concomitant with heightened statistical infarction percentage and myocardial injury markers, including CK-MB, NT-proBNP, and cTnI. The co-administration of free rhein and curcumin notably mitigated these indicators, although they remained distinguishable from baseline healthy levels. On the contrary, the administration of CurcRhe Nano, comprising rhein and curcumin, to mice resulted in the restoration of these parameters to levels akin to those in the healthy control group, underscoring CurcRhe Nano’s remarkable myocardial protective efficacy against I/R injury in mice.

**Figure 8:**
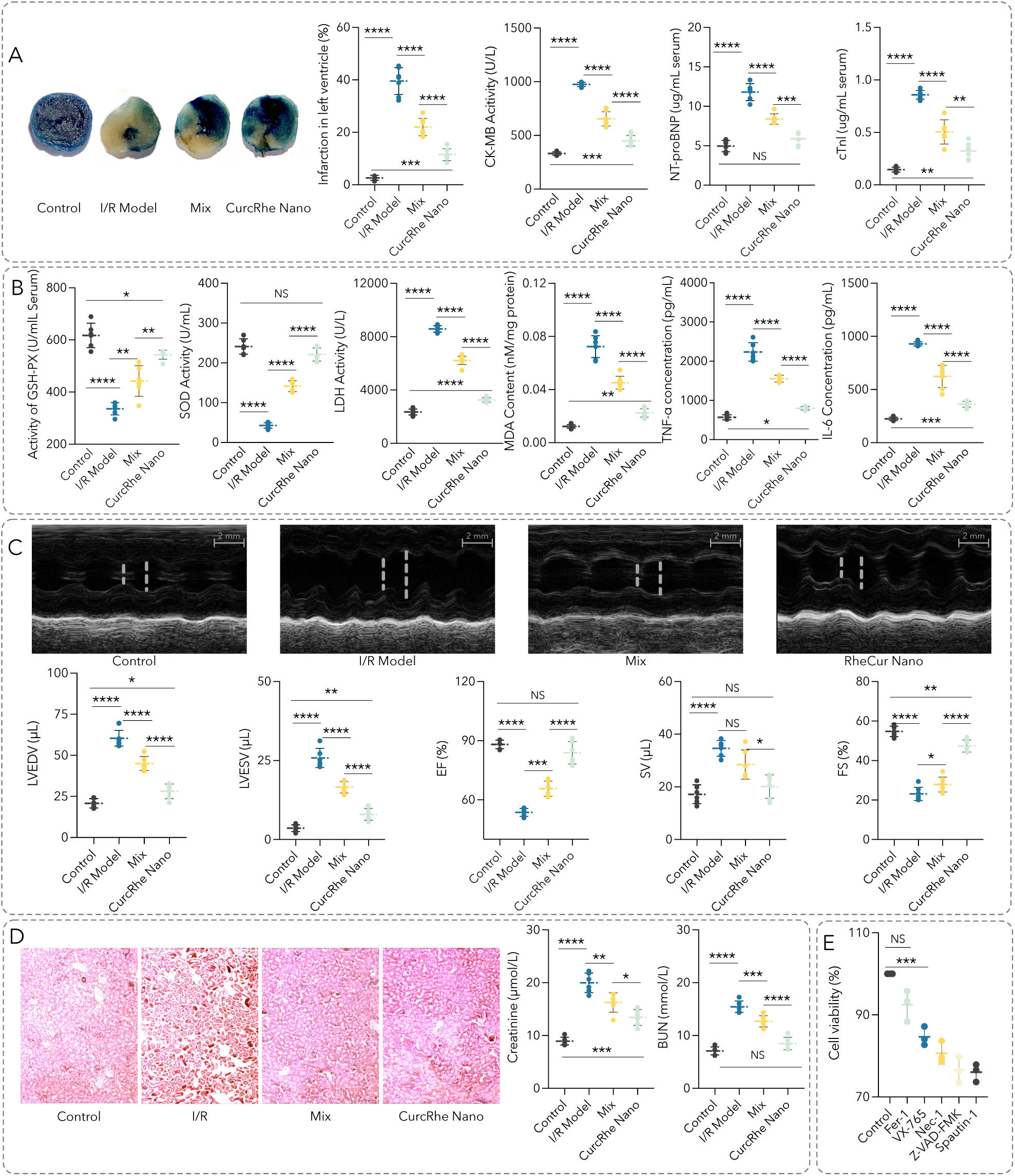
CurcRhe Nano, formulated with curcumin and rhein, protects against myocardial ischemia/reperfusion (I/R) injury in mice: (A) Reduced heart tissue infarction and normalisation of myocardial injury markers (creatine kinase myocardial band (CK-MB), N-terminal pro-hormone of brain natriuretic peptide (NT-proBNP), cardiac troponin I (cTnI)); (B) Normalised inflammatory cytokines (tumour necrosis factor-alpha (TNF-*α*), interleukin-6 (IL-6)) and oxidative stress factors (malondialdehyde (MDA), glutathione peroxidase (GSH-PX), super-oxide dismutase (SOD), lactate dehydrogenase (LDH)); (C) Normalised cardiac volumes and function (left ventricular end-diastolic volume (LVEDV), left ventricular end-systolic volume (LVESV), stroke volume (SV), ejection fraction (EF), fractional shortening (FS)); (D) Renal improvement shown by representative H&E staining of kidney sections and increased levels of blood urea nitrogen (BUN) and creatinine similar to those of the healthy control group; (E) Predominant cell death induction via ferroptosis in H9c2 cellular I/R model (Fer-1). Number of mice in each group: n = 6. * p < 0.05; ** p < 0.01; *** p < 0.001; **** p < 0.0001. H&E original magnification, 4X.

Additionally, CurcRhe Nano demonstrated outstanding anti-oxidant and anti-inflammatory activities. As shown in Figure 8B, I/R injury evidently suppressed the activities of antioxidant enzymes such as glutathione peroxidase (GSH-PX) and superoxide dismutase (SOD), while concurrently elevating lactate dehydrogenase (LDH) activity and concentrations of malondialdehyde (MDA), Tumour Necrosis Factor-alpha (TNF-*α*), and interleukin-6 (IL-6). Notably, the combination of free rhein and curcumin significantly ameliorated the levels of these markers. Moreover, the CurcRhe Nano treated group displayed reversed levels comparable to those observed in the healthy control group, once again underscoring the potential therapeutic efficacy of CurcRhe Nano in mitigating I/R-induced perturbations in oxidative stress and inflammatory markers.

Echocardiographic findings (Figure 8C) revealed notable distinctions between groups. In the I/R group, there were marked increases in left ventricular end-diastolic volume (LVEDV) and left ventricular end-systolic volume (LVESV), alongside reduced stroke volume (SV). Concomitantly, diminished ejection fraction (EF) and fractional shortening (FS) indicated a substantial decline in the left ventricular contractile capacity of mice within this group compared to the healthy control. Conversely, both the mixture group and CurcRhe Nano-treated group exhibited significant improvements in these parameters compared to the I/R group. Particularly noteworthy were the CurcRhe Nano-treated mice whose parameters were restored to levels comparable to those of the healthy control group, suggesting a compelling recuperative effect of CurcRhe Nano treatment on left ventricular function following I/R injury.

The consequential impairment caused by I/R significantly compromises kidney functionality. The kidneys’ heightened metabolic rate and reliance on continuous blood perfusion render them particularly susceptible to such injury. Episodes of reduced blood flow (ischemia) inflict harm on renal tissues due to inadequate oxygen and nutrient supply. Subsequent restoration of blood flow (reperfusion) triggers an inflammatory cascade and the generation of reactive oxygen species, amplifying the injury inflicted upon kidney tissues. Figure 8D depicted the histological examination using H&E staining of kidney sections. The images revealed evident glomerular atrophy accompanied by the loss of proximal tubular structure and vacuolisation of select tubular epithelial cells within the I/R group. Remarkably, a significant improvement in these pathologic features was observed in the treated groups, indicating a substantial improvement compared to the I/R group.

In addition, I/R injury caused a significant increase in the levels of blood urea nitrogen (BUN) and creatinine, which is indicative of renal impairment. Co-administration of free rhein and curcumin notably mitigated these elevated levels, suggesting a protective role against I/R injury. Notably, the nanoformulation comprising rhein and curcumin demonstrated an exceptional capacity to restore BUN and Creatinine levels, aligning closely with those of the healthy control group. This robust restoration underscores the promising potential of this nano-based intervention in mitigating the adverse effects of I/R injury on renal function, presenting a compelling therapeutic avenue for addressing such damage.

The cell death mechanisms associated with CurcRhe Nano were investigated (Figure 8E) using a cellular I/R model employing H9c2 cells, a continuous cell line derived from embryonic rat heart tissue. Pre-treatment of cells with Fer-1, a ferroptosis inhibitor, resulted in cell viability comparable to that of the control groups. This observation suggests that CurcRhe Nano induces cell death mainly through the mechanism of ferroptosis.

#### 3.5.3 PyBer Nano for Breast Cancer Treatment

Given the global prominence of breast cancer and the significant mortality rates associated with lung cancer^48^, a comprehensive evaluation involving breast cancer cell lines (4T-1, MCF-7, MDA-MB-231, HCC1939), lung cancer cell lines (NCIH460, A549), and normal cell lines (HK-2, NRK-52E) was conducted to assess the cytotoxicity of PyBer Nano (formulated with pyrene and berberine). Illustrated in Figure 9A, both free berberine and PyBer Nano demonstrated substantial cytotoxicity across NCI H460, A549, HCC1939, MCF-7, MDA-MB-231, 4T-1, HK-2, and NRK-52E cell lines. Notably, PyBer Nano exhibited heightened cytotoxicity against the triple-negative breast cancer (TNBC) MDA-MB-231 and breast cancer 4T-1 cells. Intriguingly, pyrene, a constituent of the formulation, exhibited no discernible cytotoxic effects at the tested concentrations across all cell types. These findings suggest that the cytotoxic impact primarily originates from berberine and the nano formulation itself.

**Figure 9:**
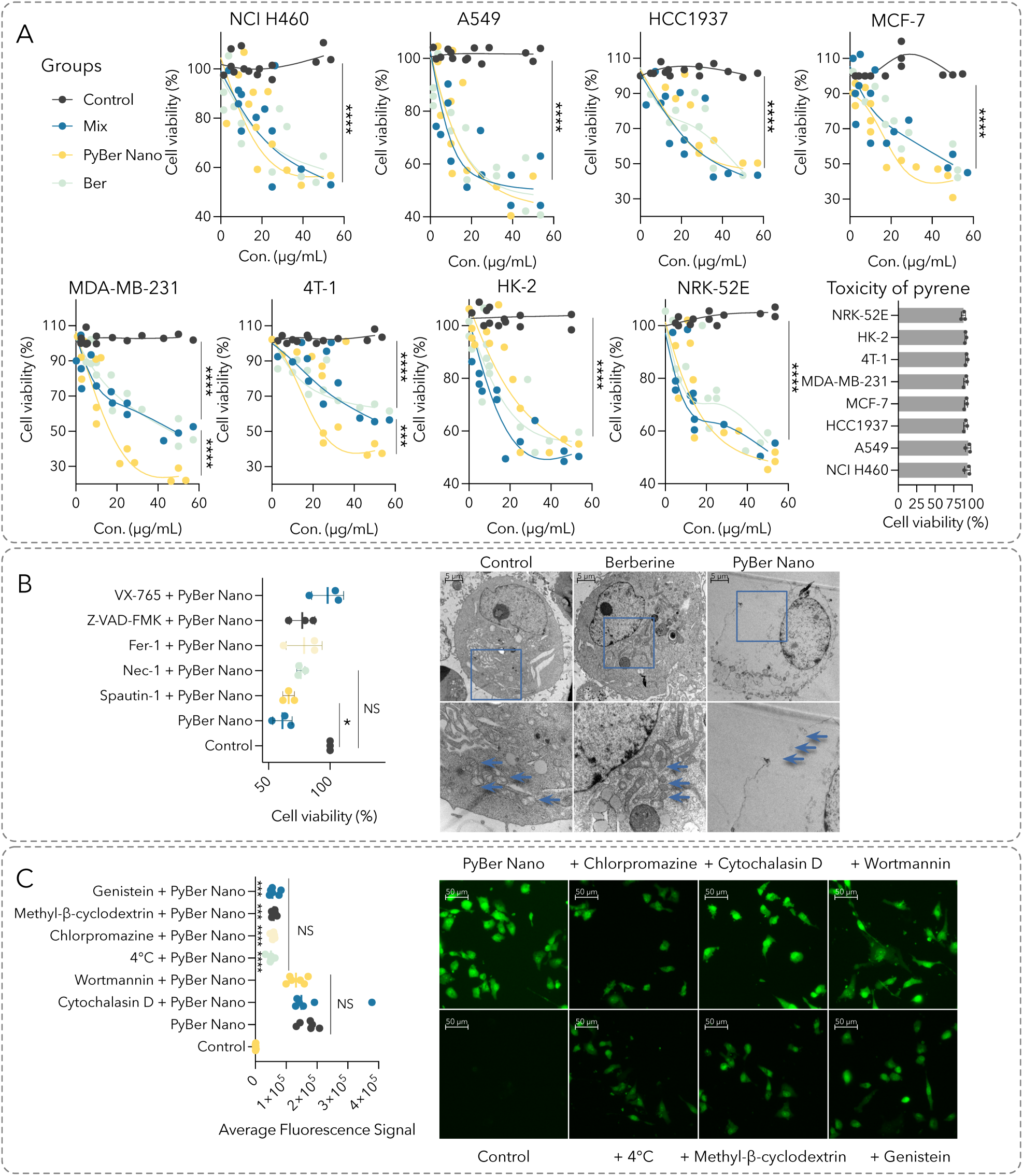
PyBer Nano, formulated with pyrene and berberine (Ber), displayed (A) heightened cytotoxicity in vitro against MDA-MB-231 cells compared to its constituent components, (B) which was achieved by instigating multiple cell death pathways, encompassing pyroptosis (VX-765), apoptosis (Z-VAD-FMK), ferroptosis (ferrostatin-1, Fer-1), and necroptosis (necrostatin-1, Nec-1), (C) through both caveolae-mediated (genistein, methyl-beta-cyclodextrin) and clathrin-mediated (chlorpromazine) endocytic pathways.* p < 0.05; ** p < 0.01; *** p < 0.001; **** p < 0.0001. Bar in TEM, 5 µM. Bar in representative fluorescence images, 50 µM.

**Figure 10:**
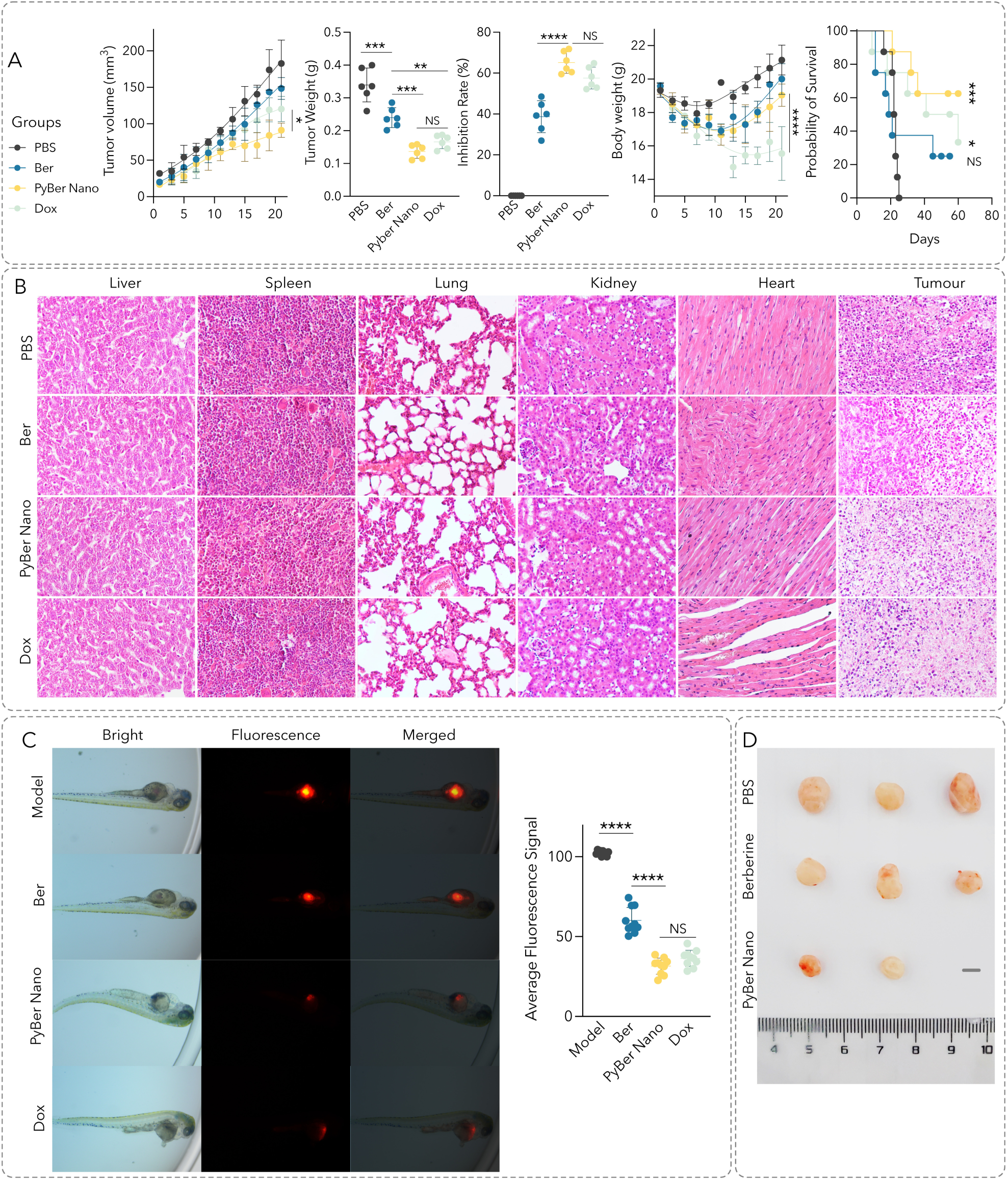
PyBer Nano, formulated with pyrene and berberine (Ber), displayed (A) Potent inhibition of MDA-MB-231 cancer comparable to doxorubicin (Dox) in mice, improved survival, and maintained bio safety without altering mouse body weight; (B) H&E staining of *ex vivo* tissue sections further confirmed the safety of PyBer Nano compared to Dox. (C) Significant suppression of tumour growth in a zebrafish model with human MDA-MB-231 cells labelled using DiI. (D) The representative *ex vivo* tumour tissues from mice subjected to various treatments. * p < 0.05; ** p < 0.01; *** p < 0.001; **** p < 0.0001. Number of mice in each group: n = 6. Number of zebrofish in each group: n = 15. Bar in TEM, 5 µM. Bar in representative fluorescence images, 50 µM. H&E original magnification, 4X.

In investigating the mechanism underlying the cytotoxic effects of PyBer Nano on MDA-MB-231 cells, a range of inhibitors targeting various cell death pathways were utilized. Figure 9B illustrates the outcomes wherein inhibitors associated with pyroptosis (VX-765), apoptosis (Z-VAD-FMK), ferroptosis (ferrostatin-1, Fer-1), and necroptosis (Necrostatin-1, Nec-1) effectively attenuated the cytotoxic effects of PyBer Nano. Notably, these inhibitors restored cell viability to levels akin to the control group. This observation strongly suggests that the cytotoxicity induced by PyBer Nano in MDA-MB-231 cells is mediated through pathways linked to pyroptosis, apoptosis, ferroptosis, and necroptosis. In addition, transmission electron microscopy (TEM) analysis revealed distinct alterations in the cells treated with PyBer Nano, characterized by a ruptured cell membrane and the absence of mitochondria and other organelles. This contrasted starkly with the intact cell membrane and typical mitochondrial morphology observed in normal cells. These observations further corroborate the multiple pathways of cell death induced by treating with PyBer Nano.

To delineate the endocytic pathways utilized by PyBer Nano, MDA-MB-231 cells were pre-treated with distinct cellular uptake inhibitors. Statistical analyses, along with representative fluorescence imaging, are detailed in Figure 9C. Notably, cells subjected to a 4°C incubation exhibited a considerable reduction in fluorescence intensity, indicative of an energy-dependent uptake mechanism for PyBer Nano. Furthermore, diminished fluorescence intensity observed in cells treated with genistein, methyl-beta-cyclodextrin, and chlorpromazine suggests a predominant utilisation of caveolae-mediated and clathrin-mediated endocytic pathways for PyBer Nano uptake. These findings provide insights into the specific cellular mechanisms (both caveolae-mediated and clathrin-mediated) governing the internalisation of PyBer Nano, elucidating fundamental aspects of its endocytic pathways.

The *in vivo* therapeutic effectiveness of PyBer Nano was investigated using nude mice implanted with MDA-MB-231-derived xenograft tumours. Statistical analyses detailing alterations in body weight, tumour volume, tumour weight, inhibition rate, and survival probability among mice treated with various formulations are depicted in Figure 9A. At the specified concentration, groups treated with doxorubicin (Dox) and PyBer Nano exhibited the most notable tumour inhibition, characterized by the smallest tumour volume and weight. However, mice treated with Dox experienced the greatest decline in body weight and significantly lower survival probability compared to those in the PyBer Nano group. The representative *ex vivo* tumour tissues from mice subjected to various treatments are depicted in Figure 9D. The representative tumour-bearing mice received different treatments are shown in Figure 3 in the Supplementary Information. These findings indicate that PyBer Nano demonstrates comparable anti-cancer efficacy to Dox against MDA-MB-231 cancer while exhibiting superior safety profiles.

Furthermore, histological analysis of *ex vivo* tissue sections using H&E staining (Figure 9B) provided additional evidence affirming the safety profile of PyBer Nano. In the Dox group, histopathological examination of cardiac tissue sections revealed substantial deviations from normal cardiac myocyte morphology. The myocardial cells displayed disorganized arrangement, noticeable vacuolisation, severe myocardial fibrosis, cellular hypertrophy, and a notable increase in apoptotic cells. Conversely, the tissues from the PyBer Nano group exhibited morphological similarities to those treated with PBS, demonstrating a comparable tissue structure to PBS-treated tissues. This similarity indicates the absence of significant adverse effects induced by PyBer Nano.

The therapeutic efficacy of PyBer Nano underwent evaluation within a zebrafish model harbouring human MDA-MB-231 tumour cells labelled with the fluorescent cell tracker DiI. The red fluorescence evident in zebrafish represented the proportional presence of MDA-MB-231 tumour cells. Remarkably, the model group showcased the most intense fluorescence and largest red area, denoting robust tumour cell proliferation. Conversely, zebrafish subjected to PyBer Nano or doxorubicin (Dox) exhibited diminished fluorescence intensity and reduced areas, indicative of substantial suppression of tumour cell proliferation. Statistical analysis of average fluorescence signal revealed that both PyBer Nano and Dox significantly attenuated intensities compared to the model group (p < 0.0001), signifying their notable therapeutic efficacy. Importantly, no statistically significant distinction emerged between PyBer Nano and Dox, affirming comparable abilities of PyBer Nano in inhibiting MDA-MB-231 tumour cells relative to doxorubicin.

#### 3.5.4 More Potential Applications

A vast landscape of potential applications unfolds in the intricate realm of Pure Nano systems, as illustrated below.

##### 3.5.4.1 NIR fluorescence imaging

NIR fluorescence imaging is a non-invasive technique that utilises visible light to stimulate fluorescent molecules, which emit NIR light for highly sensitive and specific imaging. The lower tissue scattering, absorption, and autofluorescence of NIR wavelengths in comparison to visible light enable imaging of deep tissues with higher resolution and penetration, resulting in clearer images with minimal interference from biological autofluorescence.^49^ Nonetheless, NIR fluorescence imaging poses several challenges such as non-specific binding and limited availability of imaging agents. Besides, the hydrophobic nature and low solubility of many imaging agents also limit their usefulness in imaging applications. Furthermore, ensuring the stability of these agents during storage and imaging procedures is crucial for optimal imaging efficiency and avoiding false signals.

The aforementioned Pure Nnao systems including PyRhe Nano (formulated with pyrene and rhein) and CurcRose Nano (formulated with curcumin and rose bengal) can be excited by either UV light (100 nm to 400 nm) or visible light (400 nm to 700 nm), resulting in subsequent emission of visible or NIR light beyond 900 nm. Besides, the formation of nanoparticles presents advantages such as targeted delivery capabilities, enabling these systems to fully utilise the unique features of nanoparticles. Furthermore, with the incorporation of conjugated molecules with absorption ranges that extend into the NIR region, the potential applications of nanoparticle systems can be significantly expanded, resulting in a broadening of their utilisation in various fields such as photothermal therapy, photoacoustic imaging and NIR II fluorescence imaging (ranging from 1000 nm).

##### 3.5.4.2 X-ray based imaging

In X-ray based imaging, achieving optimal contrast-to-noise ratios is crucial, and this requires a substantial difference in X-ray attenuation around the targeted tissues. Contrast agents containing elements with a high atomic number (Z) can enhance tissue differentiation due to the general increase in X-ray attenuation with atomic number. Among such agents, iodinated small molecules have been widely used as contrast agents owing to the high atomic number (Z = 53), high X-ray absorption coefficient and chemical flexibility.^50^ Iodinated small molecule contrast agents used in clinic can be classified into ionic monomers, ionic dimers, non-ionic monomers and non-ionic dimers based on their dissociation and structures. Ionic contrast agents typically have higher osmolarity, toxicity levels and anaphylactic reactions compared to non-ionic contrast agents. Non-ionic contrast agents, on the other hand, have lower osmolarity, toxicity levels, and anaphylactic reactions. Non-ionic dimers offer several advantages over non-ionic monomers, including even lower osmolarity, which reduces pain and minimises the risk of renal and cardiovascular complications.^51^ Nonetheless, non-ionic dimers have higher viscosity, which slows down excretion and may result in painful injections.

To tackle these obstacles, researchers have explored the development and evaluation of nano contrast agents by entrapping customary small molecule iodinated contrast agents into polymer-based nanoparticles. This strategy has resulted in noteworthy enhancements, such as amplified contrast enhancement, protracted circulation, appropriate osmolarity, optimal viscosity and augmented targeting capabilities.^52^ Nevertheless, these polymer-based nanoparticles tend to have a lower loading capacity, which necessitates a larger dose and higher concentration.^16^

The Pure Nano systems enable the facile formulation of iodine-containing conjugated molecules into stable and aqueous nanoparticles, showing great potential as an X-ray contrast agent owing to their exceptionally high iodine content. Notably, CurcRose Nano (formulated with curcumin and rose bengal) has been prepared with a high iodine content ratio of around 45%. Besides, the formation of nanoparticles confers these iodine-containing conjugated molecules to exploit fully the distinctive features of nanoparticles, which could enhance image quality, improve diagnostic accuracy, mitigate the systemic toxicity and ultimately lead to better patient outcomes. With further refinement of these contrast agents and increased research to confirm their effectiveness, it holds the potential for these novel nano X-ray contrast agents to emerge as a promising choice in medical imaging.

##### 3.5.4.3 Photo-based therapy

Photo-based therapy utilises distinct light sources, such as visible light and NIR, to stimulate a photosensitiser that converts the energy into other formats, including heat (photothermal therapy, PTT)^53^ and reactive oxygen species (photondynamic therapy, PDT)^54^, to effectively treat various diseases. The key advantages of photo-based therapy include its high specificity, minimal invasiveness and low toxicity. Due to their unique properties, the conjuncted molecules, such as IR-820^55,56^, pyrene^57^, rhein^58^, berberine^59^, camptothecin^60,61^ and curcumin^62^, have been formulated into various nano structures^63,64^ for exploring their potential in photodynamic therapy (PDT) and photothermal therapy (PTT) for the treatment of various ailments. The formation of Pure Nano systems of the conjuncted molecules provide them with improved nano-structural stability and aqueous dispersity as well as significantly altered photophysical properties, thus further expanding their potential applications in these fields.

## 4 Conclusion

A novel, universal approach has been developed for constructing stable and aqueous nanoparticle systems through the self-aggregation and dispersion of highly hydrophobic conjugated molecules without the use of any excipients. By classifying conjugated molecules based on their ionisation types and degrees, suitable aqueous nanoparticle systems can be formed through intensive intermolecular interactions. The contribution of preparative parameters in nanoparticle formation is negligible compared to the significant impact of formulation parameters like molecular groupings, molar ratios, organic solvents and aqueous solutions. During the spontaneous aggregation of conjugated molecules, the formation of surface charges play a crucial role in the growth and physiochemical stability of nanoparticles. This universal method circumvents solubility issues in highly hydrophobic conjugated molecules by eliminating excipients and enabling up to 100% drug loading capacity without associated toxicity risks. The *in vivo* experiments validated the remarkable efficacy of diverse Pure Nano systems in mitigating DSS-induced acute ulcerative colitis, ameliorating myocardial I/R injury, and demonstrating efficacy in mouse models of cancer. Additionally, these systems show potential for more applications such as NIR fluorescence imaging, X-ray-based imaging, and various photo-based therapies including photothermal and photodynamic therapies. Consequently, this method could revolutionise nano drug delivery systems, improve therapeutic efficacy, reduce toxicity, and ease the challenges of drug doses delivery to patients.

## Acknowledgements

We gratefully acknowledge the support of Chengdu Fuer Pharma Technology in enabling this research. Additional funding was provided by the Youth Innovation Team of Higher Education of Shandong Province (2021KJ094), the National Natural Science Foundation of China (NSFC 82003922), and China Post doctoral Science Foundation, No.13 Special Fund (2020T130334).

## Author contributions

Conceptualisation, H.X.; Methodology & Investigation, H.X., W.Z., P.H. and Q.L.; Formal Analysis, H.X. and W.Z.; Resources, H.X., Q.L. and W.Z.; Writing - Original Draft, H.X.; Writing – Review & editing, H.X., M.A., C.L. and M.F.; Visualisation, H.X.; Supervision, H.X.; Funding Acquisition, H.X., Q.L. and W.Z.

## Declaration

The authors state no conflict of interest.

