## Supplementary Information for "Pure Nano Genesis: Pioneering Universal Aqueous Nanostrategies from Pure Molecules to Revolutionise Diverse Applications"

*2024-02-15*

***Address:*.**

[1]*School of Basic Medicine, Qingdao University, Qingdao, China, 266071.*

[2]*College of Chemistry and Pharmaceutical Sciences, Qingdao Agricultural University, Qingdao, China, 266000.*

[3]*Department of Electrical Engineering, University of California, Merced, Merced, CA, 95343.*

[4]*Chemical, Biological, and Bioengineering Department, College of Engineering, North Carolina, Agricultural and Technical State University, Greensboro, NC, 27411.*

[5]*Department of Radiology, University of Michigan, Ann Arbor, MI, USA, 48109.*

[6]*Department of Biomedical Engineering, University of Michigan, Ann Arbor, MI, USA, 48109.*

[7]*Applied Physics Program, University of Michigan, Ann Arbor, MI, USA, 48109.*

† *These author contributed equally to this work.*

***Corresponding Author:***

* *Dr. Haijun Xiao, Ph.D*

**

Table of Contents 2

1 Materials 4

4 Results 17

References 19

### 1 Materials

#### 1.1 Chemical Molecules for Pure Nano Preparation

Pyrene, naphthofluorescein, nile red, amonafide, acid green 25, IR-820, rose bengal sodium, eosin Y disodium salt, methylene blue, and rhodamine B were procured from Sigma Aldrich (Shanghai, China). Pigment violet 23 was obtained from Tianfu Chemical (Henan, China), while Tanshinon IIB came from Zeye Biotechnology (Shanghai, China). Ellipticine was sourced from Aikon Chem (Jiangsu, China). Psoralen, tryptanthrin, tanshinone I, aristolochic acid, ellagic acid, hypericin, camptothecin, and anonaine were purchased from Push Biotechnology (Chengdu, China). Mitoxantrone dihydrochloride, sodium tanshinone IIA sulfonate, and topotecan hydrochloride were acquired from Selleck (China). Berberine hydrochloride, curcumin, and rhein were obtained from Zelang Medical Technology (Nanjing, China)

#### 1.2 Solvent Ingredients for Pure Nano Preparation

Glycine, hydrochloric acid, sodium acetate, acetic acid, sodium cacodylate trihydrate, citric acid, sodium citrate, sodium phosphate monobasic, sodium phosphate dibasic, sodium barbital, sodium hydroxide, acetone, acetonitrile (ACN), dimethylformamide (DMF), dimethyl sulfoxide (DMSO), ethanol (EtOH), methanol (MeOH), and tetrahydrofuran (THF) were procured from Sigma Aldrich (Shanghai, China).

#### 1.3 Pathway Inhibitors

Necrostatin-1, ferrostatin-1, Z-VAD-FMK, belmacasan (VX-765), spautin-1, wortmannin, cytochalasin D, genistein, methyl-beta-cyclodextrin were obtaeind from Selleck (China).

#### 1.4 Enzyme-Linked Immunosorbent Assay (ELISA) Kits

ELISA kits for tumour necrosis factor-alpha (TNF-$\alpha$), interleukin 6 (IL-6), interleukin 8 (IL-8), interleukin 12 (IL-12), creatine kinase myocardial band (CK-MB), cardiac troponin I (cTnI), and N-terminal prohormone of brain natriuretic peptide (NT-proBNP) were procured from Sigma Aldrich (Missouri, USA). Additionally, ELISA kits for oxidative stress factors (superoxide dismutase (SOD), glutathione peroxidase (GSH-PX), malondialdehyde (MDA), lactate dehydrogenase (LDH)), and kidney injury indicators (creatinine, blood urea nitrogen (BUN)) were supplied by Abcam (Massachusetts, USA).

##### 1.5 Others

Dextran sodium sulfate (DSS, Mw 36-50 kDa) was obtained from MP Biomedicals (California, USA). Haematoxylin & eosin (H&E) staining solution, Hoechst 33258, 2,3,5-Triphenyltetrazolium Chloride (TTC), evans blue, and DiI were purchased from Solarbio Science & Technology (Beijing, China). Dulbecco's modifed Eagle's medium (DMEM), penicillin streptomycin mixture and fetal bovine serum (FBS) were applied from ThermoFisher (Massachusetts, USA). MTT solution was obtained from Solaibio (Beijing, China).

All materials were received and stored according to instructions to maintain stability and integrity until required. Unless specified, all other materials were either prepared in-house or procured from Sigma Aldrich (Shanghai, China).

#### 2 Animals and Cells

##### 2.1 Animals

*In vivo* studies involved BALB/c-nude female mice (aged 4-6 weeks) and C57BL/6 male mice (aged 6-8 weeks) procured from Weitonglihua Experimental Animal Technology (Jinan, China). Animals underwent a week-long acclimatisation period in the laboratory prior to experimentation. The animal facility provided controlled conditions, maintaining temperature (22±2°C), humidity (50-60%), and a 12-hour light-dark cycle.

##### 2.2 Cells

Cell lines MDA-MB-231, NCI-H460, A549, HCC1937, MCF-7, 4T-1, HK-2, NRK-52E, and H9c2 were sourced from American Type Culture Collection (Maryland, USA) and cultured in Dulbecco's Modified Eagle Medium (DMEM) supplemented with 100 U/mL penicillin-streptomycin and 10% fetal bovine serum (FBS). Maintenance involved incubation at 37°C with 5% $CO_2$.

##### 2.3 Approvals and Compliance

Approval from the Animal Research Committee of Qingdao University (animal welfare assurance number: 14-0027) was obtained for all procedures, ensuring strict adherence to the Guide for the Care and Use of Laboratory Animals. Ethical protocols were strictly followed according to guidelines established by the Qingdao University Institutional Animal Care and Use Committee (IACUC), emphasising measures to minimize animal discomfort and distress throughout the study.

### 3 Methods

#### 3.1 Pure Nano Preparation and Characterisation

##### 3.1.1 Molecular Properties

Chemical information for the molecules was obtained from public databases such as ChemSpide, PubChem and DrugBank. LogP, solubility and $pK_a$ values were either sourced from literature or predicted using software tools such as ALOGPS from the Virtual Computational Chemistry Laboratory and JChem from ChemAxon. Batch prediction of molecular properties were performed based on the SMILES structure of the molecules. The solubility classification of molecules follows the United States Pharmacopeia (USP) criteria[1], which is also outlined in Table 1.

Table 1: USP solubility criteria

| Descriptive term | Part of solvent required per part of solute | solubility ($mg/mL$) |
|---|---|---|
| Very soluble | less than 1 | $> 1000$ |
| Easily soluble | from 1 to 10 | $100 - 1000$ |
| Soluble | from 10 to 30 | $33 - 100$ |
| Sparingly soluble | from 30 to 100 | $10 - 33$ |
| Slightly soluble | from 100 to 1,000 | $1 - 10$ |
| Very slightly soluble | from 1,000 to 10,000 | $0.1 - 1$ |
| Practically insoluble | more than 10,000 | $< 0.1$ |

The electrostatic surface potential of molecules were predicted using a Graph-Convolutional Deep Neural Network (ESP-DNN)[2] and visualized with the web-based molecular graphics software NGL Viewer. The colour gradient was specified to depict the potential energy ranging from -50 to 50 in units of kcal/mol, where the negative energy is denoted by shades of red and the positive by shades of blue. It is worth noting that the reported values for a particular property can vary slightly between different sources due to factors such as the method of measurement, experimental conditions and calculation protocols used.

##### 3.1.2 Nanoparticle Preparation and Optimisation

Various water-soluble organic solvents such as acetone, acetonitrile (ACN), dimethylformamide (DMF), dimethylsulfoxide (DMSO), ethanol (EtOH), methanol (MeOH), and tetrahydrofuran (THF) were used as the organic phase for the preparation of nanoparticles.

For the aqueous phase used in nanoparticle preparation, as shown in Table 2, a variety of aqueous solutions with different buffering capacities and buffering ranges were prepared.[3] Buffering range describes the $pH$ range within which a buffer can effectively maintain a stable $pH$. The $pH$ values in Table 2 correspond to the buffering range of each solution, which is dependent on the specific components added. While buffering capacities refer to the efficiency of a buffer in neutralising added acids or bases within that $pH$ range. Increasing the concentration of buffer components, such as by using higher concentration stock solutions, can enhance the buffering capacity when additional molecules are added to the solution.

Thus, it is crucial to consider both the buffer type ($pH$ range) and buffer concentration (buffering capacity) relative to the quantity of specific molecules for effective nanoparticle preparation in this unique system. Prior to nanoparticle preparation, the $pH$ of each aqueous solution was adjusted to a desired target value through the addition of hydrochloric acid or sodium hydroxide.

Table 2: Aqueous solutions with different buffering ranges

| Aqueous phase | $pH$ | $pK_a$ |
|---|---|---|
| $H_2O$ | 7.0 | 14.0 |
| Glycine HCl buffer | 2.2–3.6 | 2.35 |
| Sodium acetate buffer | 3.6–5.6 | 4.76 |
| Cacodylate buffer | 5.0–7.4 | 6.27 |
| Citrate buffer | 3.0–6.2 | 6.4 |
| Sørensen's phosphate buffer | 5.8–8.0 | 7.20 |
| Barbital buffer | 6.8–9.2 | 7.98 |
| Glycine NaOH buffer | 8.6–10.6 | 9.78 |
| Phosphate–citrate buffer | 2.2–8.0 | 7.20, 6.40 |

Pure Nano systems were generated through a rapid self-aggregation and dispersion method with slight modifications[4]. Briefly, specific amounts of small molecules were dissolved in organic solvents, forming a combined solution. This solution was then precipitated in an aqueous solution to produce a nanoparticle dispersion. Table 3 in the Manuscript delineates detailed specifications for each formulation, including molecular grouping, mass quantities, molecular ratios, designated organic solvent as the oil phase, $pH$ levels of the aqueous phase, and respective volumes of the aqueous phase. Nanoparticle powder was obtained *via* meticulous lyophilisation, a freeze-drying technique preserving structural integrity and stability. The resultant product was stored under light-protected conditions at 4°C to prevent degradation until further use.

Various process parameters were investigated to understand their impact on nanoparticle preparation, including the dissolving of small molecules, the self dispersion approach, the speed and duration of stirring, as well as the inclusion or exclusion of organic solvents in the suspensions. In addition, the formulation of nanoparticles was also evaluated by examining the influence of small molecule ratios, organic solvents and aqueous phases on the nanoparticle formation. All the resulting nanoparticles were evaluated according to their particle size, surface charge and size distributions.

##### 3.1.3 Nanoparticle Characterisation

The particle sizes, distributions and surface charges of nanoparticles in dispersions were measured by a Malvern Zetasizer Nano. Around one millilitre of nanoparticle dispersion was added into a plastic cuvette and the particle sizes were measured after two minutes of equilibration. The surface charges of nanoparticles were also measured by injecting a nanoparticle dispersion into a disposable folded capillary cell.

The morphologies of nanoparticles were observed by a scanning electron microscope (SEM). The nanoparticle dispersion was dropped onto an aluminium foil and left in a fume hood until the moisture evaporated completely to obtain a dried product. The nanoparticles were then coated with a layer of gold powder to increase the electrical conductivity before being investigated on a scanning electron microscope (NanoSEM 450).

The investigation into the fluorescence behaviour of nanoparticles or small molecules in various solvents was conducted using an FLS 1000 photoluminescence spectrometer under controlled conditions with a set temperature of 25°C.

The assessment of nanoparticle stability encompassed a comprehensive analysis, evaluating changes in nanoparticle sizes, distributions, and surface charges. This examination involved controlled dilution with appropriate solvents, storage conditions, and variations in environmental pH values. Periodic measurements were conducted to monitor alterations in nanoparticle physicochemical properties, providing insights into their stability across diverse environmental conditions.

#### 3.2 Acute ulcerative colitis (UC) Induction in Mice & Therapeutic Assessment

##### 3.2.1 DSS-induced UC in Mouse Model

The induction of ulcerative colitis (UC) in a mouse model[5] involved the dissolution of dextran sulfate sodium (DSS) in distilled water to formulate a 2.5% (W/W) DSS solution. C57BL/6 male mice weighing between 18-22 g were randomly assigned to four groups: a healthy control group receiving regular drinking water, a UC group exposed to 2% DSS for 7 days, a free rhein group administered 100 mg/kg once daily for 7 days via intragastric administration (ig), wherein rhein was suspended in distilled water with sodium carboxymethyl cellulose, and a PyRhe Nano (formulated with pyrene and rhein) group receiving the same dose and administration route for 7 days.

Monitoring throughout the experiment encompassed the recording of body weight changes. Subsequent to the experimental period, both blood samples and colonic tissues were harvested for further analysis.

###### 3.2.2 Determination of Fecal Water Content

To determine fecal water content in mice, on the 6th day of the experimental protocol, mice were individually housed in metabolic cages equipped with wire mesh flooring for 24 hours to facilitate fecal collection.

Following the collection period, freshly collected fecal samples underwent immediate weighing to determine their initial weight. Subsequently, the samples were subjected to desiccation or lyophilisation to obtain a consistent dry weight. The calculated difference between the initial weight and the dry weight allowed for the precise quantification of fecal water content, providing crucial insights into colonic water absorption dynamics. The following formula was used to calculate the fecal water content[6]:

$$\text{Fecal water content (\%)} = \left(\frac{\text{Fresh feces weight} - \text{Dry feces weight}}{\text{Fresh feces weight}}\right) \times 100\%$$

###### 3.2.3 Quantification of Serum Pro-inflammatory Cytokines *via* ELISA

The quantification of serum pro-inflammatory cytokines TNF-$\alpha$ , IL-6, IL-8, and IL-12 employed the enzyme-linked immunosorbent assay (ELISA) technique.[7] ELISA, a highly sensitive and specific immunological assay, facilitates the precise measurement of target proteins or analytes in biological samples. Specifically designed ELISA kits for TNF-$\alpha$ , IL-6, IL-8, and IL-12 detection were utilised in this study.

The ELISA process involves immobilizing target-specific capture antibodies onto a solid surface, typically a microplate well. Subsequent incubation of serum samples allows the binding of target cytokines to the immobilized antibodies. After rigorous washing to eliminate unbound components, detection antibodies conjugated with enzymes are introduced, forming antibody-antigen complexes. Addition of a substrate for the enzyme generates a quantifiable colorimetric or fluorescent signal, directly correlating to the concentration of the target cytokines present in the serum samples. Precise measurements were obtained using a microplate reader for accurate cytokine level assessment.

###### 3.2.4 Determination of Colon Lengths

Post-experiment, the determination of colon lengths in mice served as a crucial assessment of colonic integrity.[8] Upon animal sacrifice, meticulous dissection was performed to expose the gastrointestinal tract. Delicate excision of the colon was carried out to avoid undue stretching or manipulation that might influence length measurements.

The isolated colons were carefully positioned, and precise measurements from the cecum to the rectum were obtained using calibrated instruments, ensuring the natural curvature of the colon was maintained. Skilled personnel conducted these measurements to guarantee accuracy and consistency. Any observed anomalies or structural deviations in the colon were meticulously recorded for comprehensive evaluation.

##### 3.2.5 Histological Analysis

Histological analysis of colonic tissues involved the preparation of 5 μm-thick paraffin-embedded tissue sections. Initial tissue fixation in formalin followed by embedding in paraffin wax facilitated the microtome-based sectioning to acquire the desired tissue thickness.

Deparaffinisation and rehydration of the tissue sections were carried out sequentially using alcohol gradients. Subsequent staining with haematoxylin & eosin (H&E) followed standardised protocols, enabling selective staining of cell nuclei in blue and cytoplasmic components in pink, respectively. The stained sections underwent dehydration, xylene clearance, and were coverslipped using mounting media.

Light microscopy with appropriate magnification and imaging software facilitated the observation and capture of detailed images, enabling meticulous examination of tissue morphology and any potential pathological features evident in the colonic tissue sections.

#### 3.3 Ischemia/Reperfusion (I/R) Injury Modeling and Therapeutic Evaluation in Mice

##### 3.3.1 I/R mice Model Construction and Treatment

C57BL/6 male mice were divided randomly into four groups, each comprising 6 mice: a healthy Control group receiving standard animal treatment, an I/R group subjected to ischemia/reperfusion (I/R) injury through a 45-minute occlusion of the left anterior descending coronary artery with non-absorbable nylon sutures followed by 24 hours of reperfusion, a Mix group administered a compound mixture (curcumin and rhein) at a dose of 20 mg/kg *via* intragastric administration for 1 hour prior to I/R, and a CurcRhe Nano (formulated with curcumin and rhein) group at a dose of 10 mg/kg *via* intravenous administration for 1 hour before I/R.

Intragastric administration was selected for the Mix group due to the considerable limitations posed by the notably low water solubility of curcumin, a hydrophobic compound. This inherent property of curcumin renders its intravenous (IV) administration infeasible.

Post-experiment, blood samples were extracted from the abdominal aortic artery for further analysis. Additionally, hearts and kidneys were isolated for detailed investigations aimed at evaluating specific markers and parameters associated with I/R injury.

##### 3.3.2 Transthoracic Echocardiography

Transthoracic echocardiography, a non-invasive imaging modality, was conducted using a high-resolution ultrasound imaging system Vevo2100 (VisualSonics, Canada), coupled with a 30 MHz mechanical transducer. This imaging technique enabled high-resolution cardiac assessments in mice. Mice were anaesthetized with 2% isoflurane in 100% oxygen and placed on a warming platform to maintain optimal body temperature during imaging procedures.

The imaging protocol involved two-dimensional guided M-mode echocardiography in the parasternal long-axis view. This approach facilitated the precise measurement of crucial cardiac parameters such as ejection fraction (EF), fractional shortening (FS), left ventricular end-diastolic volume (LVEDV), and left ventricular end-systolic volume (LVESV), essential markers for assessing cardiac function and performance.

Data analysis was based on the averaging of 3–6 cardiac cycles from a minimum of two scans per mouse, ensuring robust and reliable measurements of cardiac parameters while minimizing variations inherent in cardiac cycles and imaging sessions.

##### 3.3.3 TTC and Evans Blue Double-Staining

The application of Evans Blue dye and 2,3,5-triphenyltetrazolium chloride (TTC) enables the distinct identification of myocardial regions based on differential colour changes induced by these dyes.[9] Following a 24-hour reperfusion period in C57BL/6 male mice subjected to cardiac I/R injury, a 2.0% Evans Blue dye solution was intravenously administered *via* the jugular vein. This dye selectively stains non-ischemic tissue regions within the myocardium, resulting in a vivid blue colouration that distinctly delineates areas unaffected by ischemic damage, particularly evident in the non-ischemic left ventricle (LV).

Subsequent to Evans Blue staining, the excised heart sections, each approximately 1 mm thick, were immersed in a 1.0% TTC solution (Sigma Aldrich, US). TTC interacts with metabolically active cells, causing viable myocardial tissue to adopt a red hue due to the formation of formazan. Consequently, regions hosting active cellular metabolism exhibit a noticeable reddish tint, indicating non-infarcted myocardium.

In contrast, infarcted tissue remains unstained and appears white after TTC incubation. The absence of TTC-induced red colouration highlights areas devoid of metabolic activity, facilitating clear demarcation of infarcted regions against the blue-stained non-ischemic tissue backdrop. This differential colouration enables precise differentiation between ischemic, non-ischemic, and infarcted areas within the myocardium. The distinct colours resulting from Evans Blue and TTC staining facilitate accurate localization and assessment of myocardial injury, enabling quantitative analysis and detailed mapping of the extent of tissue damage.

Digital imaging of stained sections was analysed via computerized planimetry using ImageJ software. This approach precisely outlined and quantified the infarct (unstained) and non-ischemic left ventricle (red-stained) areas, yielding the infarct size as a percentage of the total LV area. This method provided a quantitative measure of myocardial damage ensuing from I/R injury.

##### 3.3.4 Histological Analysis

Upon collection, the kidney tissues were fixed in 10% formaldehyde for preservation. Subsequent paraffin embedding facilitated the creation of thin tissue sections, approximately 5-10 micrometers thick, suitable for microscopic analysis.

Following deparaffinisation and rehydration, these sections were stained with H&E dyes, highlighting cellular components. Haematology imparted a blue-purple hue to cellular nuclei, whereas eosin provided contrast by staining cytoplasm and extracellular structures pink or red.

Post-staining, the sections were readied for microscopic examination to discern intricate tissue morphology and cellular details using a light microscope.

##### 3.3.5 Assessment of Blood Biomarkers

Serum markers, encompassing cardiac injury indexes (CK-MB, cTnI, NT-proBNP), inflammatory cytokines (TNF-α, IL-6), oxidative stress factors (SOD, GSH-PX, MDA, LDH), and kidney injury indicators (creatinine, BUN), underwent meticulous evaluation using specialised assay methodologies. Techniques employed varied, including enzymatic, immunological, and spectrophotometric assays tailored to assess specific biomolecules.

##### 3.3.6 Simulating *in Vitro* Myocardial Ischemia/Reperfusion (I/R) Injury using Anoxia/Reoxygenation (A/R) Model

To replicate myocardial ischemia/reperfusion (I/R) injury in vitro, H9c2 cells at approximately 70% confluency were subjected to a controlled sequence of hypoxic and reoxygenation phases.[10] Initially, these cells were cultured in glucose- and serum-depleted Dulbecco's Modified Eagle Medium (DMEM) within a specialized hypoxia chamber. This chamber, rendered airtight, was supplied with a gas mixture of 95% nitrogen ($N_2$) and 5% carbon dioxide ($CO_2$) to induce a hypoxic state at a regulated temperature of 37°C for 45 minutes, mimicking the ischemic phase.

Subsequent replacement of the culture medium with standard DMEM and cultivation of cells under typical cell culture conditions at 37°C with 5% $CO_2$ for 24 hours simulated the reoxygenation or reperfusion phase following ischemia.

In pursuit of delineating the distinct cell death pathways activated by CurcRhe Nano within an I/R injury model, H9c2 cells were subjected to pre-treatment with a range of death pathway inhibitors. These included Fer-1 (a ferroptosis inhibitor), VX-765 (a pyroptosis inhibitor), Nec-1 (a necroptosis inhibitor), Z-VAD-FMK (an apoptosis inhibitor), or Spautin-1 (an autophagy inhibitor). The aim was to modulate or inhibit specific cell death mechanisms triggered subsequent to CurcRhe Nano exposure.

Following a 24-hour incubation period post-exposure to CurcRhe Nano, the impact of the inhibitor pre-treatment on cell viability was assessed using the MTT assay. Briefly, cells were incubated with CurcRhe Nano for 24 hours, then treated with MTT solution for an additional 4 hours. Formazan crystal formation was quantified by measuring absorbance at 570 nm using a microplate reader (BIOTEK, ELX-800, USA).

#### 3.4 Cancer Modeling and Therapeutic Assessment in Mice

##### 3.4.1 *In Vitro* Cytotoxicity Measurement

The MTT (3-(4,5-dimethylthiazol-2-yl)-2,5-diphenyltetrazolium bromide) assay, a foundational method for assessing cellular viability and toxicity, was utilised to investigate the impact of Berberine (Ber), PyBer Nano (formulated with pyrene and berberine), and a physical mixture of pyrene and berberine across an array of cell lines including MDA-MB-231, NCI-H460, A549, HCC1937, MCF-7, 4T-1, HK-2, and NRK-52E.

In a controlled experimental setup, these diverse cell lines were seeded into 96-well plates and subjected to varying concentrations (5, 10, 25, 50 μg/mL) of the aforementioned treatments. Post-incubation, the introduction of MTT solution facilitated formazan crystal formation proportional to viable cell numbers. Measurement of the resulting absorbance provided insights into the relative cytotoxicity induced by each treatment across the spectrum of cell types.

Moreover, the cytotoxicity of pyrene was evaluated at its maximum concentration corresponding to the content in the formulation for comparative analysis.

##### 3.4.2 Pathways of Cell Death

In elucidating the specific cell death pathways activated by PyBer Nano, MDA-MB-231 cells were subjected to pre-treatment with a range of death pathway inhibitors. These included Fer-1 (a ferroptosis inhibitor), VX-765 (a pyroptosis inhibitor), Nec-1 (a necroptosis inhibitor), Z-VAD-FMK (an apoptosis inhibitor), or Spautin-1 (an autophagy inhibitor).

Subsequent to a 1-hour pre-treatment with these inhibitors, the cells were incubated for an additional 24 hours to ascertain the effects on cell viability post-PyBer Nano exposure. Following this incubation period, 10 μL of MTT solution was introduced to the cells and incubated for an additional 4 hours. Measurement of the resultant formazan crystals' absorbance enabled an assessment of the activated cell death pathways in response to PyBer Nano exposure.

Besides, the samples of PyBer Nano-treated MDA-MB-231 cells were meticulously prepared for scrutiny under a Transmission Electron Microscope (Hitachi 800, Tokyo, Japan). Following a gentle centrifugation process, the supernatant was meticulously removed, leaving behind a pellet sized akin to a large grain of rice. This prepared sample facilitated a detailed investigation into cellular morphology to discern potential manifestations indicative of diverse cell death pathways.

##### 3.4.3 Uptake Pathways: Cellular Uptake Routes

To unravel the cellular uptake pathways engaged by PyBer Nano within MDA-MB-231 cells, a series of inhibitors including chlorpromazine, genistein, methyl-beta-cyclodextrin, wortmannin and cytochalasin D were applied to disrupt specific cellular uptake mechanisms. This comprehensive approach aimed to decipher the primary pathways involved in PyBer Nano internalisation within the cellular milieu.

Initial treatment of MDA-MB-231 cells with these inhibitors occurred at a reduced temperature of 4°C for 0.5 hours, strategically impeding distinct cellular uptake processes. Subsequent incubation with PyBer Nano at a concentration of 20 μg/mL for an additional 2 hours allowed observation of the nanoparticles' internalisation within the cells.

Post-treatment, the removal of excess or unbound nanoparticles from the culture dishes facilitated the observation of nanoparticle uptake patterns. Employing a confocal laser scanning microscope, the fluorescence intensity attributed directly to the individual components constituting PyBer Nano within the cells was recorded. This analysis aimed to delineate the predominant cellular uptake pathways utilised by PyBer Nano within the MDA-MB-231 cells under the influence of various inhibitors.

##### 3.4.4 Establishment of Subcutaneous Tumours in Nude Mice

BALB/c-nude female mice, aged 4-6 weeks, underwent a week-long acclimation before being randomly divided into four groups, each comprising 6 mice: PBS, Berberine hydrochloride (Ber), PyBer Nano and doxorubicin hydrochloride (Dox) groups.

Following the acclimation phase, mice were subcutaneously inoculated with 0.1 mL of MDA-MB-231 cell suspension.[11] Tumours emerged at the inoculation site within 5 days, and upon reaching a diameter exceeding 5 mm, indicating successful model establishment, treatment ensued. Intravenous administrations of Dox (3 mg/kg, solution), Ber (10 mg/kg, solution), or PyBer Nano (10 mg/kg, dispersion) occurred every other day up to Day 21.

Regular monitoring of tumour volume and body weight at 3-day intervals was conducted. Tumour volume estimation was determined utilising a mathematical formula[12] derived from measurements of the longest and shortest diameters of the tumour ($r^2 \geqslant 0.97$ for nude mice). This approach considers the irregular shape of tumours and calculates their volume as an approximation of ellipsoids. utilising the longest and shortest diameter measurements, quantification of the tumour's size in cubic millimetres was achieved, providing a standardized method for assessing tumour growth and response to treatments.

$$\text{Tumour Volume (mm}^3\text{)} = \frac{\text{Longest Diameter} \times \text{Shortest Diameter}^2}{2}$$

Upon conclusion, mice were euthanised, and tissues comprising tumours, liver, spleen, lung, kidney, and heart underwent sectioning and Haematoxylin and Eosin (H&E) staining for histological analysis. Accurate documentation of tumour weight, inhibition rate and survival probability was carried out to assess treatment efficacy.

##### 3.4.5 *In Vivo* Distribution

The *in vivo* biodistribution of PyBer Nano post intravenous administration in BALB/c-nude female mice bearing MDA-MB-231 tumours was monitored using the In Vivo Imaging System (IVIS) within a 24-hour time frame. The IVIS system, equipped to capture specific fluorescence signals emitted by PyBer Nano components, facilitated real-time tracking of nanoparticle distribution within the live subjects.

Subsequent *ex vivo* analysis involved harvesting organs, including tumours, liver, spleen, lung, kidney, and heart at 6 and 24-hour intervals post-injection, followed by their examination under the IVIS. This comprehensive approach allowed for detailed scrutiny of the distribution patterns of PyBer Nano among various organs post-administration.

##### 3.4.6 Human Tumour Cell Transplantation in Zebrafish and Quantification of Tumour Growth

Human MDA-MB-231 tumour cells were labelled with the fluorescent cell tracker DiI[13] (1 μg/mL) and subsequently injected into the coelomic cavity of zebrafish[14], each containing 2 nL and around 100 cells. Following cell transplantation, the aquarium temperature was adjusted to 32°C, optimising the engraftment and growth of the human tumour cells within the zebrafish over 48 hours.

A total of 40 zebrafish were randomly distributed into four experimental groups, including the Model group, Berberine hydrochloride (Ber) group, PyBer Nano group, and Doxorubicin hydrochloride (Dox) group, comprising 10 animals per group. Each group received the corresponding treatment: namely Ber (50 μg/kg), PyBer Nano (50 μg/kg), or Dox (30 μg/kg), administered *via* the aquarium water for 48 hours.

Subsequent to the treatment duration, the zebrafish were anaesthetised using fishese (0.1 mg/mL) for imaging. Images were captured to assess the success of xenotransplantation, specifically focusing on the fluorescence area as an indicator of engraftment efficiency and tumour growth within the zebrafish model.

This method enables the non-invasive observation and quantification of human tumour cell growth in the zebrafish, providing a valuable platform for studying tumour behaviour and evaluating therapeutic responses *in vivo*.

#### 3.5 Statistical Analysis

The *in vivo* data underwent Analysis of Variance (ANOVA) comparisons using GraphPad Prism version 9.4.1. This method facilitated comprehensive group comparisons for statistical significance.

### 4 Results

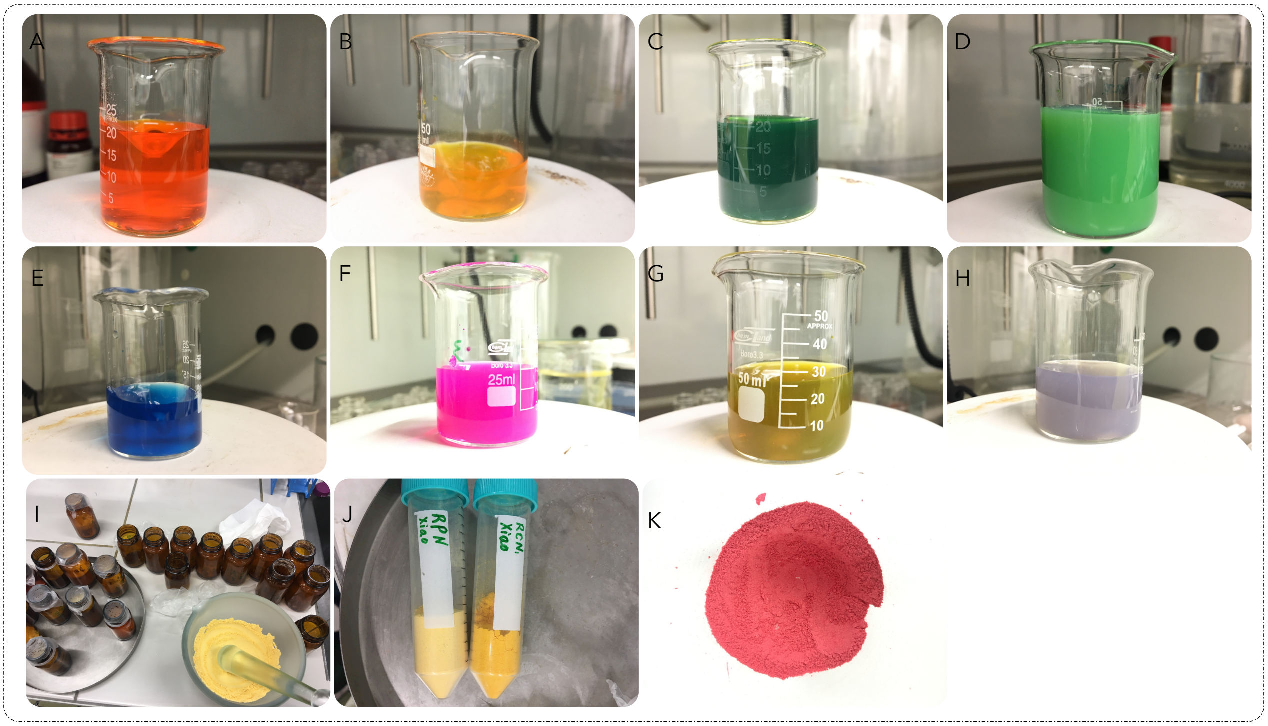

Figure 1: The representative dispersions of Pure Nano systems. (A) CurcRose (formulated with curcumin and rose bengal); (B) CurcRhe (formulated with curcumin and rhein); (C) CurcMitox (formulated with mitoxantrone dihydrochloride and curcumin); (D) PyR820 (formulated with IR820 and pyrene); (E) PyreBlue (formulated with methylene blue and pyrene); (F) PyRhod B (formulated with Rhodamine B and pyrene); (G) HyperCur (formulated with hypericin and curcumin); (H) PyHyper (formulated with hypericin and pyrene); (I) Collection of lyophilised Pure Nano powder; (J) The lyophilised powder of PyRhe and CurcRhe; (K) The lyophilised powder of CurcRose.

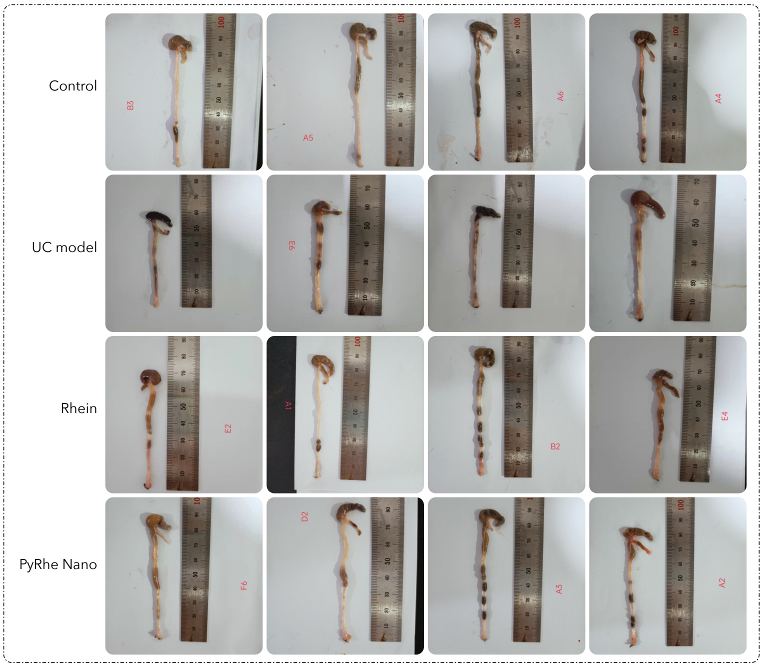

Figure 2: The representative ex vivo mouse colon tissues received different treatments.

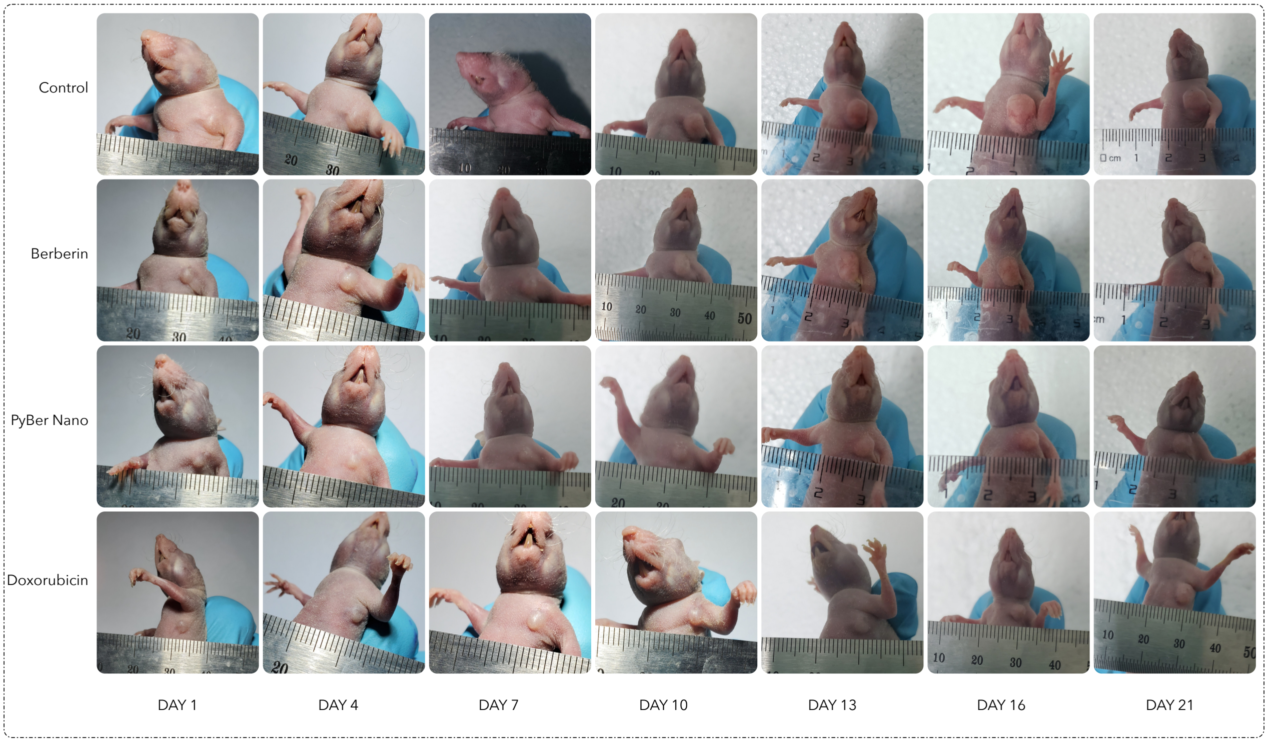

Figure 3: The representative tumour-bearing mice received different treatments.
